# Vimentin Intermediate Filaments Can Enhance or Abate Active Cellular Forces in a Microenvironmental Stiffness-Dependent Manner

**DOI:** 10.1101/2022.04.02.486829

**Authors:** Farid Alisafaei, Kalpana Mandal, Maxx Swoger, Haiqian Yang, Ming Guo, Paul A Janmey, Alison E Patteson, Vivek B. Shenoy

## Abstract

The mechanical properties of cells are largely determined by the cytoskeleton, which is a complex network of interconnected biopolymers consisting of actin filaments, microtubules, and intermediate filaments. While disruption of the actin filament and microtubule networks is known to decrease and increase cell-generated forces, respectively, the effect of intermediate filaments on cellular forces is not well understood. Using a combination of theoretical modeling and experiments, we show that disruption of vimentin intermediate filaments can either increase or decrease cell-generated forces, depending on microenvironment stiffness, reconciling seemingly opposite results in the literature. On the one hand, vimentin is involved in the transmission of actomyosin-based tensile forces to the matrix and therefore enhances traction forces. On the other hand, vimentin reinforces microtubules and their stability under compression, thus promoting the role of microtubules in suppressing cellular traction forces. We show that the competition between these two opposing effects of vimentin is regulated by the microenvironment stiffness. For low matrix stiffness, the force-transmitting role of vimentin dominates over their microtubule-reinforcing role and therefore vimentin increases traction forces. At high matrix stiffness, vimentin decreases traction forces as the microtubule-reinforcing role of vimentin becomes more important with increasing matrix stiffness. Our theory reconciles seemingly disparate experimental observations on the role of vimentin in active cellular forces and provides a unified description of stiffness-dependent chemo-mechanical regulation of cell contractility by vimentin.

**Significance:** Vimentin is a marker of the epithelial to mesenchymal transition which takes place during important biological processes including embryogenesis, metastasis, tumorigenesis, fibrosis, and wound healing. While the roles of the actin and microtubule networks in the transmission of cellular forces to the extracellular matrix are known, it is not clear how vimentin intermediate filaments impact cellular forces. Here, we show that vimentin impacts cellular forces in a matrix stiffness-dependent manner. Disruption of vimentin in cells on soft matrices reduces cellular forces, while it increases cellular forces in cells on stiff matrices. Given that cellular forces are central to both physiological and pathological processes, our study has broad implications for understanding the effect of vimentin on cellular forces in different microenvironments.

## Introduction

Cells in the body experience physical stresses in the form of tension, compression, and shear during different physical activities including muscle contraction, breathing, blood flow, body movement, rest, and sleep (*1*). It is now well known that these forces play important roles in many biological processes including tissue and organ morphogenesis, proliferation, differentiation, and gene expression (*1, 2*). More than a century ago, it was proposed that in addition to these “external” forces, non-muscle cells generate “internal” forces and can transmit these forces to their surrounding extracellular matrices (ECMs) through focal adhesions (*3*). It was later shown that these internally generated contractile forces enable adherent cells to sense and respond to the mechanical properties of their ECM. For example, cells in both two-dimensional (2D) and three-dimensional (3D) environments sense the stiffness of the ECM (*4, 5*) and generate higher traction forces when they are exposed to stiffer environments (*6–8*). Through these internally-generated contractile forces, cells also engage in a mechanical crosstalk with their ECMs as these forces can themselves reorganize and alter the local mechanical properties of the ECMs (*8, 9*).

In non-muscle cells, the internal forces are generated by the actomyosin machinery. The myosin head domains bind to actin filaments and pull on them to generate internal contractile forces which are transmitted to the ECM. The key role of myosin and actin in the generation and transmission of internal forces has been demonstrated in various studies where inhibition of either myosin motors or actin polymerization significantly decreases traction forces (*10*). Unlike actin filaments, which experience tension and for which their disruption markedly decreases traction forces, microtubules undergo compressive deformations (*11–13*) and their disruption leads to an increase in cellular contractility (*14*) and traction forces (*15*). These increases in cellular contractility and force generation have been attributed to the mechanical resistance of microtubules against contraction (*11, 12*), and activation of GEF-H1 and the Rho-Rock pathway upon microtubule depolymerization (*14, 16*).

While the roles of the actin and microtubule networks in the generation and transmission of internal forces are known, it is not clear if and how intermediate filaments impact cellular forces. For example, experimental studies in the literature on fibroblasts, which are the most common cell type in connective tissues, report three different scenarios; upon depletion of vimentin intermediate filaments in fibroblasts, cell tractions forces have been observed (i) to decrease (*17–19*), (ii) to increase (*20–22*), or (iii) to remain the same (*23*). This discrepancy calls into question the role of vimentin intermediate filaments in the transmission of internally generated contractile forces to the ECM.

To answer this question, we introduce a chemo-mechanical cell model that accounts for all the key cellular components involved in the generation and transmission of cellular forces to the ECM. Using this model along with experimental data, we first show that vimentin intermediate filaments (VIFs) can have opposite effects on cellular forces as they interact with the actomyosin or microtubule networks. We next demonstrate that these opposite effects of intermediate filaments compete in an ECM stiffness-dependent manner. As a result, depending on ECM stiffness, cell traction forces can increase, decrease, or remain the same upon depletion of intermediate filaments, reconciling the seemingly opposite results in the literature. To validate the predictions of our model, we studied mouse embryonic fibroblast cells derived from wild-type (VIF +/+) and vimentin-null (VIF −/−) mice on substrates with different stiffness. Our experimental results show that VIF −/− cells exhibit lower spreading area, and traction forces on soft substrates, while on stiff substrates they spread more and generate higher traction forces compared with VIF +/+ cells, in agreement with the predictions of our theory.

## Results

### Modeling the chemo-mechanical crosstalk between vimentin intermediate filaments, microtubules, and actin filaments

The cytoskeleton is composed of three interconnected biopolymers: actin filaments, microtubules, and intermediate filaments. There is growing evidence that these three biopolymeric networks physically interact with each other and this physical interaction plays an important role in many cellular functions including cell migration and polarization (*24*). To understand the role of intermediate filaments in the transmission of cellular forces to the ECM, we develop a 3D chemo-mechanical model that accounts for all the key cellular components involved in the generation and transmission of forces (see Materials and Methods). In this model, the cell cytoskeleton is treated as a continuum of representative volume elements (RVEs), each of which is comprised of (i) the myosin molecular motors, (ii) the microtubule network, (iii) the actin filament network, and (iv) the intermediate filament network (Figure 1A and Table S1). All these elements are initially uniform (independent of spatial location) and isotropic (independent of direction) as these elements have the same initial density everywhere in the cytoplasm with no preferential alignment. Starting with the uniform and isotropic conditions, we will describe later how our simulations predict nonuniform and anisotropic reorganizations of each of these cytoskeletal components and how their reorganizations contribute to the transmission of cellular forces to the extracellular environment. In what follows, we first describe how actin, myosin, and microtubules in our model respond to ECM mechanics through signaling pathways in agreement with experimental results in the literature. We then add VIFs to the model and describe how the model predicts the existence of two sets of VIFs with distinct functions required to consistently explain our experimental results.

**Figure 1.**
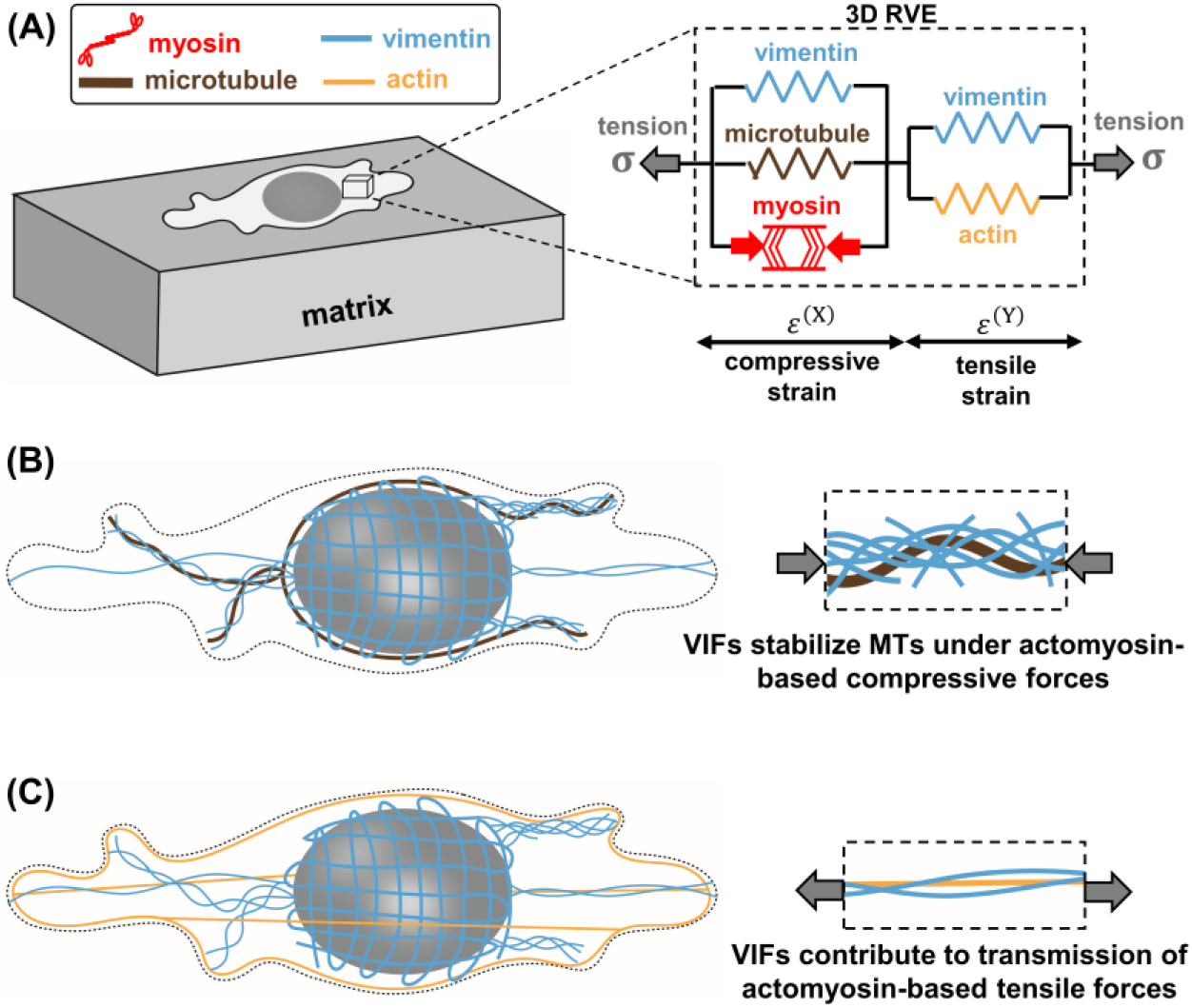
Mechanical crosstalk between cytoskeletal components. (A) The cell is modeled as a continuum of representative volume elements (RVEs), each of which is constructed by connecting (i) an active force-generating contractile element representing myosin motors, (ii) the actin filament network, (iii) the microtubule network, and (iv) two elements of the vimentin intermediate filament network. (B) The first vimentin element laterally reinforces and stabilizes microtubules under contractility-based compressive forces. (C) The second vimentin element interacts with actin filaments and is involved in the transmission of contractility-based tensile forces to the matrix.

#### Modeling the active cellular contraction

The first component of the model is myosin which generates cell internal contractility as reported experimentally (*25*) (Figures 1A and S1). We treat the average density of phosphorylated myosin motors as a symmetric tensor, *ρ*_*ij*_, whose components represent cell contractility in different directions (see SI Section 1.1) (*26*). The cell contractility *ρ*_*ij*_ generates compressive stress 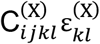 and tensile stress σ_*ij*_ in the cytoskeletal components that are in compression (e.g., microtubules) and tension (e.g., actin elements), respectively,

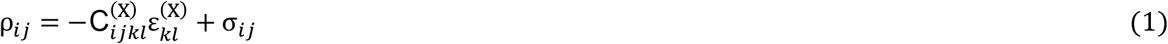

where 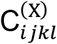 and 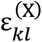 are the stiffness and strain tensors of the cytoskeletal components that are in compression (Figure 1A).

#### Modeling the adjustment of myosin phosphorylation level and force generation in response to mechanical signals

As tension is generated in the cytoskeleton (either by the intrinsic cell contractility or external tensile forces), cells increase their contractility through phosphorylation of more myosin motors which in turn generates higher cytoskeletal tension. This reciprocal mechanism is controlled by tension-activated signaling pathways such as the Rho-Rock and the Ca^2+^ pathways (Figure S1) (*27–31*). To include the signaling pathways and capture this feedback mechanism, we assume that the average of contractility in all three directions, 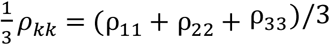, increases with the average of tension in the cytoskeleton, 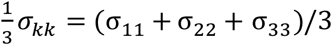,

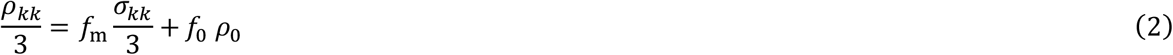

where this stress-dependent feedback mechanism is regulated by the feedback parameter *f*_m_. In the absence of tension (*σ*_*kk*_ = 0), *f*_0_ regulates the mean contractility 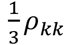 with *ρ*_0_ being the basal cell contractility (see SI Section 1.1 for *f*_m_ and *f*_0_). We have previously shown that this positive feedback between contractility and cytoskeletal tension (Figure S2) consistently explains cell responses to mechanical signals such as matrix stiffness, cell spreading area, and external forces (*32*). For example, as a result of the feedback mechanism, our simulations show that the cell contractility *ρ* and the cell-generated tensile stress *σ* increase with matrix stiffness and then reach a plateau (Figure S3), consistent with experimental observations (*33*).

#### Modeling the polymerization of the actin network in response to tension

The second component of the model is actin filaments which are connected to the myosin element in series (Figure 1A). Subsequently, the actin element experiences tension and transmits myosin-generated tensile forces to the extracellular matrix through focal adhesions as observed in experiments (*33–35*). Starting with a uniform and isotropic distribution, the stiffness of the actin network 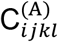 increases in proportion and in the directions of the tensile principal components of the stress tensor σ_*ij*_,

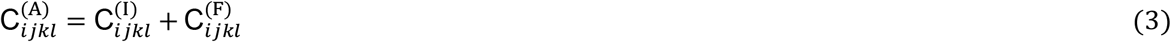

where **C**^(I)^ and **C**^(F)^ are the initial and stiffening parts of the stiffness tensor **C**^(A)^, respectively. Note that **C**^(F)^ ≈ 0 for low tensile stress σ_*ij*_ and increases with tension (but not in compression), representing the formation of actin filaments and stress fibers in response to tension as observed experimentally (*32, 36, 37*) (Figure S4). (see SI Sections 1.1 and 1.2). As a result of this stiffening and concomitant with phosphorylation of more myosin motors (Figure S5), the model shows that the cytoskeletal stiffness *C* increases with matrix stiffness and then reaches a plateau (Figure S6) in agreement with our previous experiments (*38, 39*). On the other hand, disruption of actin filaments, by setting the stiffness of the actin element *C* ^(A)^ ≈ 0, leads to decreases in cell contractility, cell-generated traction force, and cell-induced matrix deformation, consistent with experimental measurements (*40*) (Figure S7).

#### Modeling the increase in cell contractility in response to microtubule depolymerisation

The third component of the model is microtubules, which are placed in parallel with the contractile element (Figure 1A). As a result, microtubules in the model experience compression which is consistent with experimental observations (*11, 12*). Depolymerization of microtubules reduces their mechanical resistance against cell contraction and activates GEF-H1 and the Rho-Rock pathway in the model as observed experimentally (*11, 12, 14, 16*) (Figure S8, see SI Section 1.1). As a result, simulations of microtubule depolymerization show that cells become more contractile, and subsequently generate higher traction forces and stretch the matrix more (Figure S7), which are all consistent with experimental measurements (*14, 15, 41–43*).

#### Addition of vimentin intermediate filaments to the model

We next study how intermediate filaments should be connected to other cytoskeletal components in the model. To this end, we focus on fibroblasts which are relatively high contractile cells, and we study the effect of vimentin intermediate filaments on their contractile forces. Vimentin filaments, directly and indirectly, interact with both microtubule and actomyosin networks (see “Discussion and Conclusions” for experimental evidence) (*24, 44–48*). We, therefore, include two vimentin elements in the model to account for the interaction of vimentin with both microtubule and actomyosin networks (Figures 1A-1C). Using the model, we first study the effect of each of these vimentin elements, and their overall effect on cellular forces. We then experimentally validate the model predictions for fibroblasts cultured on different matrix stiffness.

#### Modeling actin-vimentin interactions

In addition to non-physical interactions through biochemical signaling, intermediate filaments also interact with actin filaments through physical contact mediated by cross-linkers, direct binding, and steric effects (*24*) as we have previously shown in our in vitro studies (*44*). A large portion of VIFs that interact with contractile actin filaments are expected to experience tensile stresses (Figure 1C) (*49*). We thus add an element of intermediate filaments in parallel with the actin element, thereby experiencing tensile stresses (Figure 1 A). The element represents the actomyosin-associated VIFs and promotes the transmission of contractility-based tensile forces to the ECM (Figure 2A, top panel). As expected, our simulations show that disruption of this element leads to decreases in the cell contractility *ρ*, the cell-generated stress *σ*, and the matrix strain *ε*_m_ (Figure S9).

**Figure 2.**
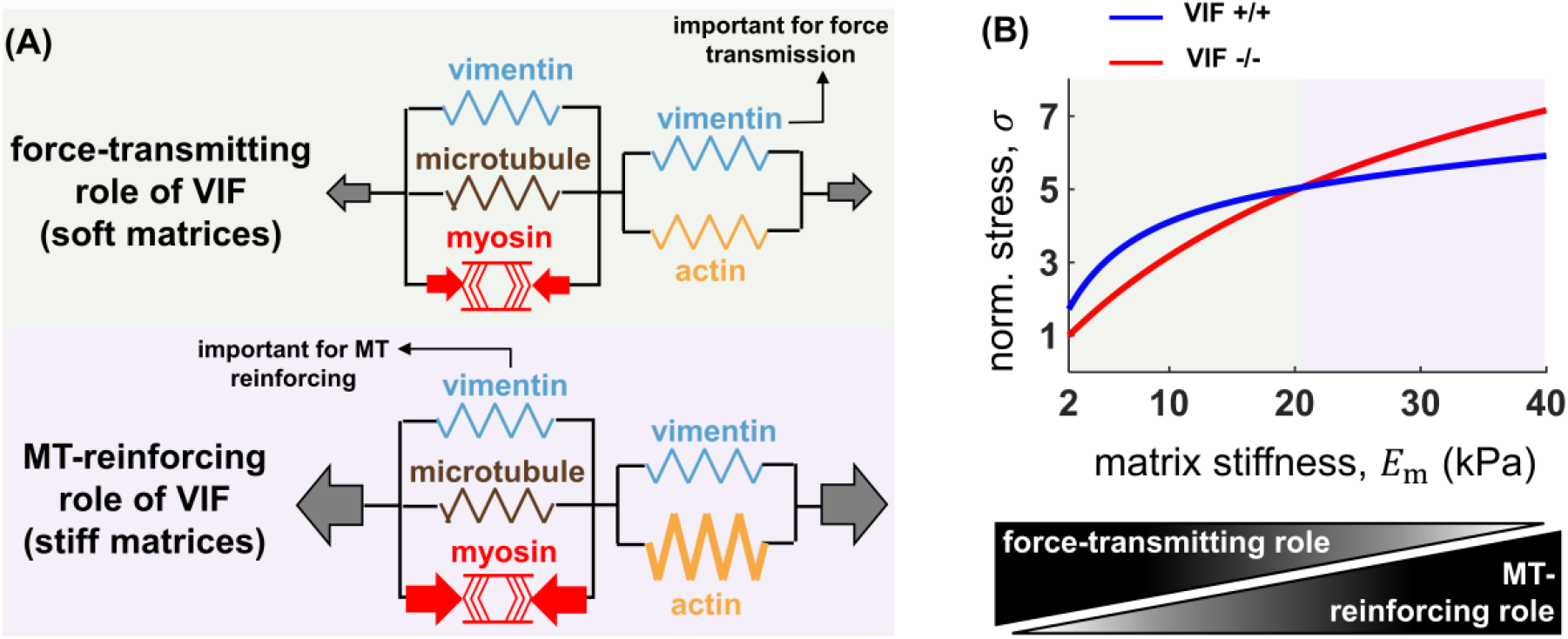
Disruption of the vimentin network can decrease or increase cell contractility depending on microenvironment stiffness (model predictions). (A) On the one hand, vimentin intermediate filaments (VIF) are involved in the transmission of actomyosin-based tensile forces to the matrix and therefore disruption vimentin can decrease cell contractility. On the other hand, vimentin filaments reinforce microtubules under compression. As a result, disruption of vimentin can lead to destabilization of microtubules, which can in turn increase cell contractility. Cells on soft substrates are characterized by a weak actomyosin network and vimentin plays an important role in the transmission of forces to the substrate (force-transmitting role of VIF), while on stiff substrates, microtubules experience high compression, and they require reinforcement from the vimentin network to withstand the compression (MT-reinforcing role of VIF). (B) The model shows how the two opposing effects of vimentin compete in a matrix stiffness-dependent manner. The model predicts that for low matrix stiffness (low cellular contractility), the force-transmitting role of vimentin overpowers its microtubule-reinforcing role and therefore disruption of vimentin decreases cellular contractility and tension. In contrast, for high matrix stiffness (high cellular contractility), disruption of vimentin increases contractility and tension as the microtubule-reinforcing role of vimentin becomes more important with increasing matrix stiffness.

#### Modeling microtubule-vimentin interactions

Microtubules interpenetrate a dense network of intermediate filaments which laterally reinforces and stabilizes microtubules under contractility-based compressive stresses (Figure 1B) (*13, 50*). Therefore, we add another element of intermediate filaments to the model in parallel with the microtubule element (Figure 1A), thereby supporting microtubules under contractility-based compressive stresses and preventing them from destabilization (Figure 2A, bottom panel). This VIF element negatively affects cellular forces by resisting cell contraction and reinforcing microtubules. Our simulations show that disruption of the VIF element in parallel with the microtubule element increases the cell contractility *ρ*, the cell-generated stress *σ*, and the matrix strain *ε*_m_ (Figure S9).

Taken together, VIFs exhibit opposite effects on cell contractility in our model. On the one hand, disruption of vimentin decreases traction force due to the force-transmitting role of VIFs, while on the other hand, disruption of vimentin increases traction force due to the microtubule-reinforcing role of VIFs (Figure 2A). We next study how these two opposite effects of vimentin compete with each other, and we examine whether this competition depends on matrix stiffness.

### Vimentin filaments can increase or decrease active cellular forces depending on matrix stiffness

With the two VIF elements in the model, we simulate the disruption of vimentin filaments to study how lack of VIFs affects cellular forces at different matrix stiffness. To this end, we set the stiffness of both VIF elements equal to zero in our model. The model predicts that at low matrix stiffness (or when actomyosin contractility is low), the force-transmitting role of vimentin filaments is more significant compared with their microtubule-reinforcing role and therefore disruption of vimentin decreases the cell contractility *ρ*, the cell-generated stress *σ*, and the matrix strain *ε*_m_ (Figures 2B and S10). In contrast, the model shows that for cells on stiff substrates (cells with high levels of actomyosin contractility), the microtubule-reinforcing role of vimentin filaments dominates over their force-transmitting role and therefore disruption of vimentin increases *ρ, σ*, and *ε*_m_ (Figures 2B and S10).

The model shows that the reorganization of the cytoskeleton with matrix stiffness regulates the matrix stiffness-dependent effect of vimentin on cellular forces. Vimentin filaments are simultaneously involved in the transmission of tensile forces to the matrix (force-transmitting role) and the reinforcement of microtubules under compression (microtubule-reinforcing role). In cells on soft substrates, actin filaments form a weak network and microtubules experience low compression. Therefore, the force-transmitting role of vimentin is more important than its microtubule-reinforcing role. As a result, depletion of VIFs in cells on soft substrates decreases cellular forces. In contrast, in cells on stiff matrices, actin filaments form a strong contractile network to transmit tensile forces to the matrix and microtubules experience high compression. Thus, the microtubule-reinforcing role of vimentin is more important than its force-transmitting role. As a result, depletion of VIFs in cells on stiff substrates causes more buckling and instability of microtubules which increase cellular forces.

### Experimental validation of model predictions: vimentin impacts cellular forces in a matrix-dependent manner

As illustrated in Figure 2B, the model predicts that disruption of vimentin has opposite effects on the cellular force at different matrix stiffness, resulting in a crossover in VIF +/+ and VIF −/− curves. The model also shows that the matrix stiffness at which the crossover occurs depends on the physical properties of the vimentin filaments network (Figure S11). For example, intermediate filaments are known to exhibit strain stiffening behavior (*51–53*), and the model predicts that tension or compression stiffening of intermediate filaments can shift the crossover point to higher or lower matrix stiffness, respectively (Figure S12). Therefore, to validate the model prediction and to examine whether the crossover occurs at a physically measurable and physiologically relevant matrix stiffness range, we culture wild-type (VIF +/+) and vimentin-null (VIF −/−) mouse embryonic fibroblast (mEF) cells on fibronectin-coated hydrogel substrates with different stiffness.

#### Matrix stiffness-dependent effect of vimentin on cell area

We have previously shown that spreading areas of both VIF +/+ and VIF −/− cells increase with substrate stiffness (*54*). However, compared with cells containing vimentin (VIF +/+ cells), our results showed that VIF −/− cells spread less on soft substrates (0.75 and 6 kPa), whereas they spread significantly more on rigid substrates (Figure 3C). These results show that vimentin can increase or decrease cell spreading area in a matrix stiffness-dependent manner. Since the cell spreading area is positively correlated with the cell traction force (*7, 55*), these experimental results may indicate that vimentin impacts cellular forces in a matrix stiffness-dependent manner, thereby supporting the predictions of our model (see also Figure S13 for our previous theoretical and experimental studies on the effect of cell spreading area on actomyosin contractility (*32*)).

**Figure 3.**
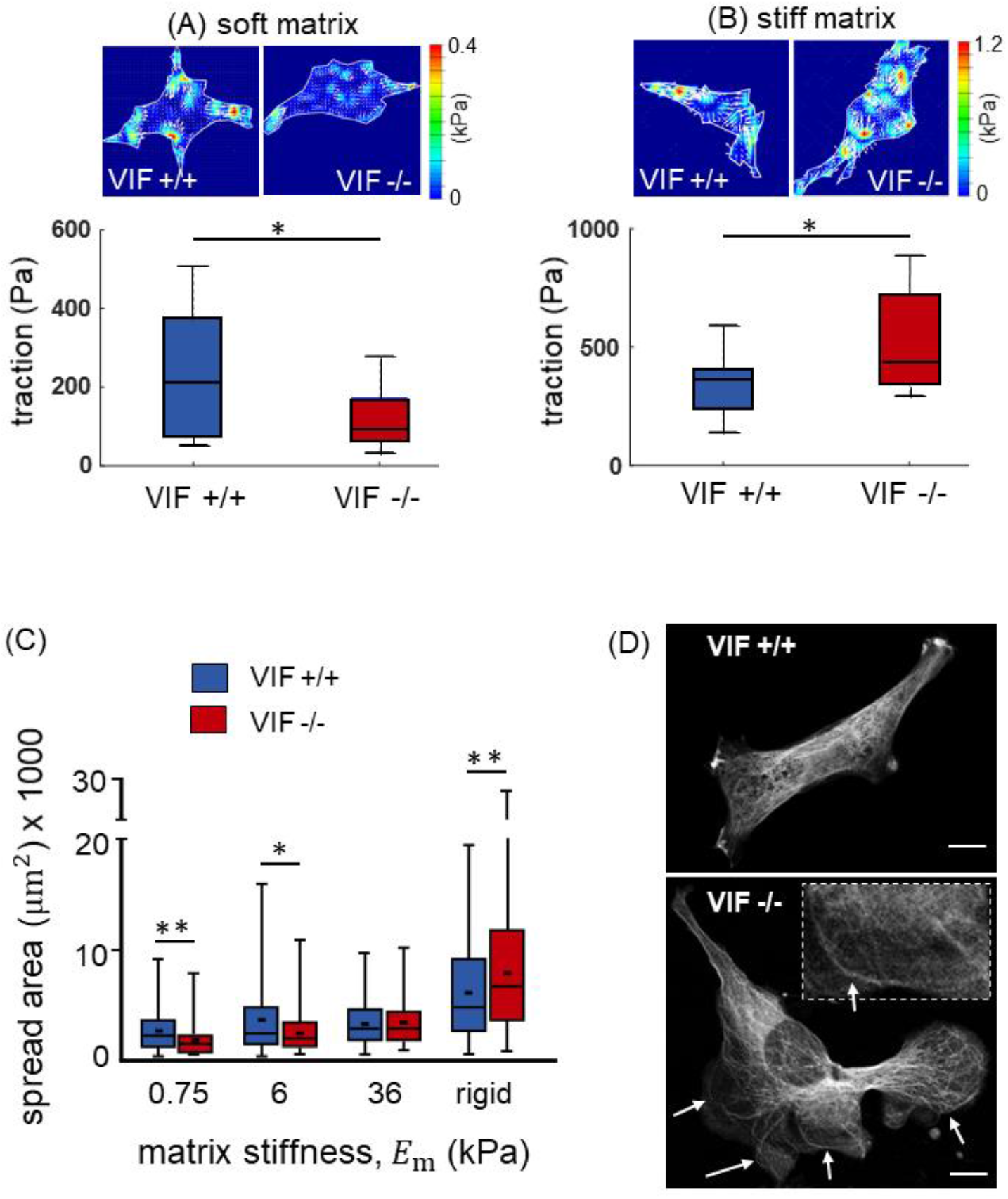
Matrix-stiffness dependent effect of vimentin on traction forces and spreading area (experimental validations). (A) Compare with control cells, VIF −/− cells generate lower traction forces on soft substrates (elastic modulus = 15 kPa and *n* = 21-27). (B) In contrast, VIF −/− cells generate higher forces on stiff substrates (elastic modulus = 40 kPa and *n* = 8-17). (C) Similarly, VIF −/− cells spread less on soft substrates, whereas they spread significantly more on rigid substrates (all *n* > 100). (D) On stiff substrates, the microtubule-reinforcing role of vimentin becomes more important as microtubules show instability and buckle abnormally (denoted by the white arrows) without the support of vimentin in VIF −/− cells (scale bar: 10 μm). error bars: standard error

#### Matrix stiffness-dependent effect of vimentin on cell traction forces

Next, we perform traction force microscopy (TFM) experiments to directly measure cellular forces of VIF +/+ and VIF −/− cells on a relatively stiff substrate. Our TFM results showthat the traction stress generated by VIF −/− cells is higher than that of VIF +/+ cells on the stiff substrate (Figures 3B and S14). Furthermore, our experimental results show that, compared with wild-type fibroblasts, microtubules in VIF −/− cells exhibit lower stability and buckle at much larger wavelengths resulting in abnormal microtubule shapes under contractility-based compressive stresses (Figure 3D). These are both in agreement with the model prediction that the microtubule-reinforcing role of VIFs becomes more important on stiff substrates, and therefore disruption of VIFs leads to instability of microtubules and an increase in cell traction force (Figure 2).

We next measure traction forces on the soft substrate and we find that in contrast to the TFM results on the stiff substrate, VIF −/− cells generate lower traction stress than that of VIF +/+ cells on the soft substrate (Figure 3A), validating the model prediction that VIFs impact traction forces in a matrix stiffness-dependent manner.

Taken together, our experiments show that vimentin impacts cell traction force and spreading area of fibroblasts in a matrix stiffness-dependent manner. Note that these results are also consistent with TFM experiments of plectin −/− fibroblasts. Plectin is a cytolinker protein that links vimentin filaments to actin filaments, myosin filaments, and microtubules, forming an interconnected network of biopolymers (*47*). In agreement with model prediction and similar to the case of VIF −/− cells on soft substrates, plectin −/− fibroblasts generate lower contractile forces on soft substrates of 4 and 8 kPa compared with wild-type cells (*56*). In contrast, on rigid substrates, plectin −/− fibroblasts exhibit a significant augmentation of actin stress fibers and focal adhesions compared with wild-type fibroblast cells (*57*). All these results show that VIFs interact with the other cytoskeletal components in a matrix stiffness-dependent manner. As a result, disruption of this interaction, by depletion of either vimentin or plectin, can have matrix-dependent effects on cellular forces.

### Vimentin intermediate filaments contribute to the long-range propagation of local forces in the cytoplasm

To further study the role of vimentin intermediate filaments in the propagation of forces, we next use optical tweezer microscopy where we generate a local force field in the cytoplasm and measure how the generated force propagates in the cytoplasm with and without the presence of vimentin filaments (*58*). In our optical tweezer microscopy experiments, a 1 μm radius (*r*) bead is dragged in the cytoplasm over 200 nm (*u*_0_) while the resulting displacement and strain fields generated around the bead in the cytoplasm are measured by visualizing the movement of surrounding fluorescently labeled mitochondria (Figure 4). As the bead moves in the cytoplasm, compressive (negative strain) and tensile (positive strain) fields are generated in the front and back of the bead, respectively. Experimental results show that both compressive and tensile fields extend significantly farther in wild-type fibroblasts than in VIF −/− cells. This is better shown in Figure 4A where we plot the local cytoplasmic displacement along the drag direction, *u*, as a function of the distance to the bead, *x*. These experiments show that the displacement field both in the front and back of the bead extends farther in VIF +/+ cells. Our simulations demonstrate that this long-range displacement propagation is due to the strain-stiffening property of vimentin both in tension and compression. As vimentin filaments undergo compressive and tensile deformations in the front and back of the moving bead, respectively, they stiffen and exhibit higher resistance against deformation (Figure S15) which in turn leads to long-range propagation of the deformation and strain fields (Figure 4). Furthermore, vimentin also protects the other cytoskeletal components under loading, as lack of vimentin in VIF −/− cells causes cytoskeletal damage in optical tweezer experiments which in turn leads to strain-softening and fast decay of the displacement field in VIF −/− cells. Together, our results show that vimentin filaments not only are involved in the propagation of local forces in the cytoskeleton, but they also increase the range of force propagation.

**Figure 4.**
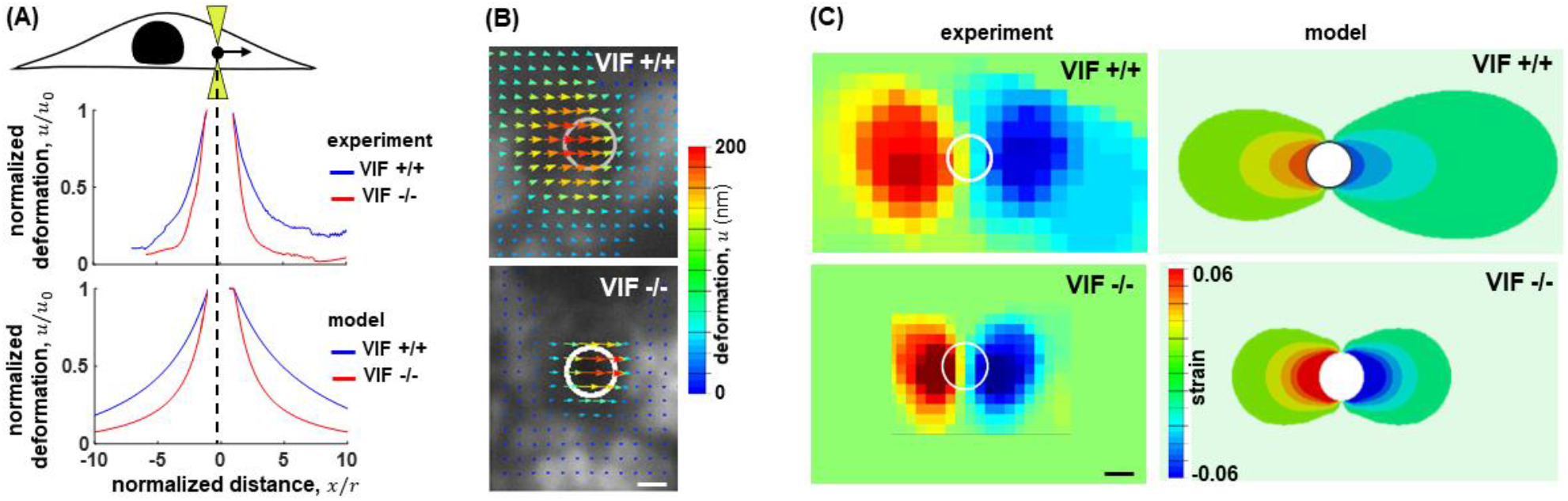
The effect of vimentin on the propagation of local forces in the cytoplasm. The involvement of vimentin in the propagation of both tensile and compressive forces is further illustrated in optical tweezer experiments where a 1-μm radius bead is dragged in the cytoplasm over 200 nm. Visualizing the movement of surrounding fluorescently labeled mitochondria shows the displacement and strain fields around the bead. Both deformation and strain fields significantly extend farther in wild-type cells than in the VIF −/− cells, indicating that vimentin plays an important role in the propagation of local forces. (Scale bars: 2 μm)

### The state of mechanical stress in the cytoskeleton determines the spatial distribution of vimentin filaments

Since the model has two VIF elements interacting with the actin and microtubule networks, we next ask how mechanical stresses are spatially distributed in each of these VIF elements. Note that as the myosin element generates contractile forces, the elements in parallel with the myosin element experience compression and resist contraction, while the elements in series are subject to tensile stresses and promote the transmission of tensile forces to the ECM (Figure 5A). Our three-dimensional model enables us to measure the stress that each of these elements experiences in different directions. Note that cells can experience different stress levels in different directions and therefore stress should be considered as a direction-dependent quantity. Thus, as described in SI Section 1, we first determine the stress tensors for the compressive (parallel) 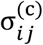 and tensile (series) 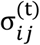 elements. We then calculate the minimum eigenvalue (minimum principal value) of the stress tensor 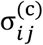 to determine the maximum compressive stress that the compressive elements experience at any point in the cytoskeleton. Similarly, we calculate the maximum eigenvalue of 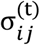 to determine the maximum tensile stress that the tensile elements spatially experience (Figure 5A). We finally plot the maximum compressive stress and the maximum tensile stress as vectors, *σ*^(c)^ and *σ*^(t)^, whose orientations represent the maximum compression direction (purple lines in Figure 5A) and the maximum tension direction (green lines in Figure 5A) in the parallel and series elements, respectively. Our simulations show that around the nucleus and in the direction perpendicular to the nuclear envelope, *σ*^(c)^ is significantly high. This reveals that parallel VIFs experience high levels of compression in the juxtanuclear region and therefore this family of VIFs is expected to appear more wavy, compressed, and buckled around the nucleus. On the other hand, *σ*^(t)^ in the basal plane is high close to the cell periphery, indicating that the series VIFs undergo high tensile stresses in the basal plane close to the cell boundary. Therefore, this family of VIFs is predicted to appear as aligned and stretched fibers close to the periphery of the cell.

**Figure 5.**
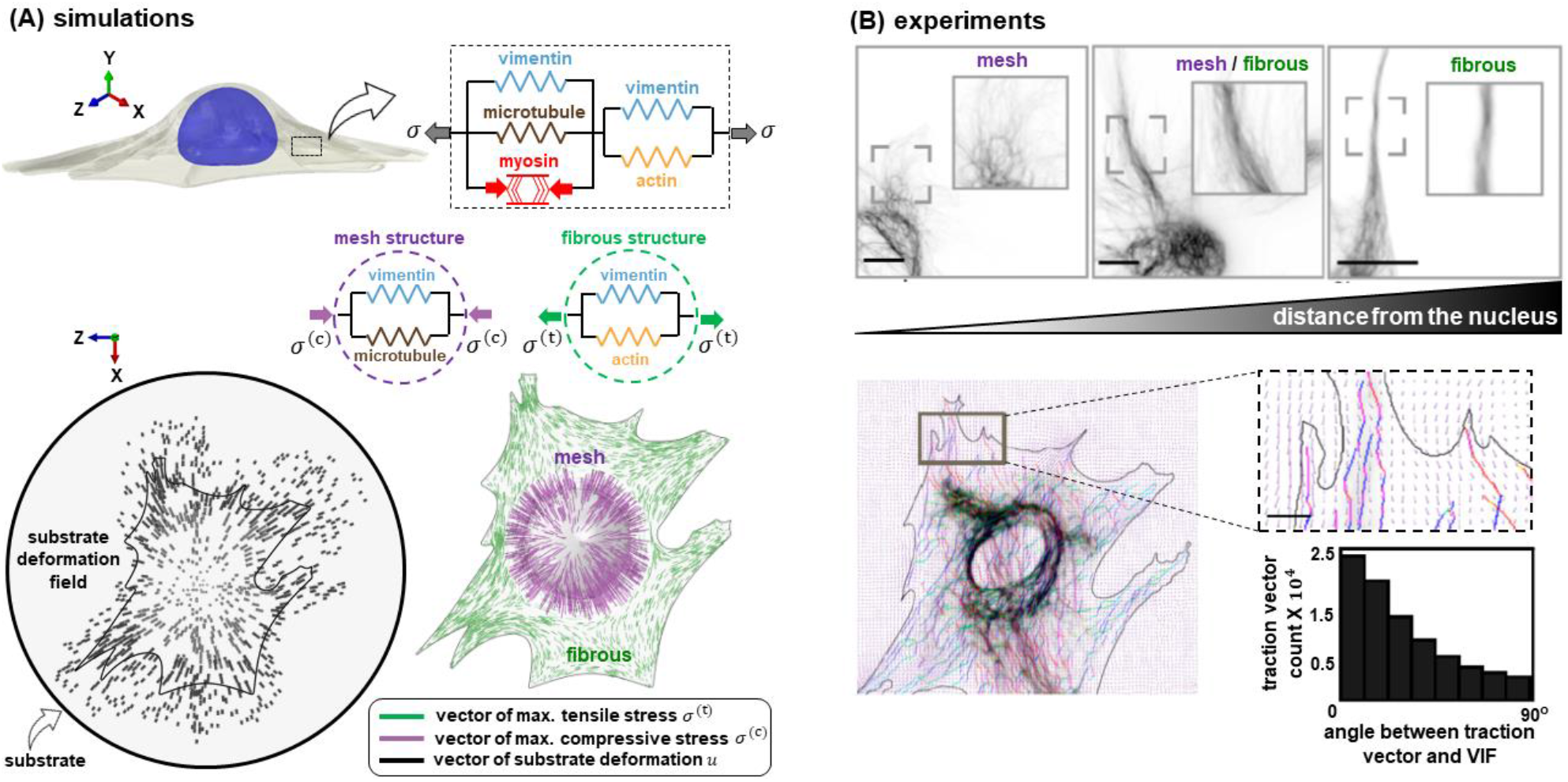
Distribution of vimentin filaments is determined by the state of mechanical stress in the cytoskeleton. (A) A fibroblast is simulated on a deformable substrate as a 3D continuum of RVEs, each of which is composed of five elements. As the active element (representing myosin motors) generates internal contractile forces, the elements in parallel with the active element experience compression, while the elements in series undergo tensile stresses. We determine the maximum compressive stress *σ* ^(c)^ and the maximum tensile stress *σ* ^(t)^ that the compressive and tensile elements experience, respectively, at each RVE. Our simulations show high compressive stress *σ* ^(c)^ around the nucleus in the direction perpendicular to the nuclear envelope, and high tensile stress *σ* ^(t)^ in the basal plane close to the cell periphery. Thus, the juxtanuclear vimentin filaments are expected to appear wavy, forming a mesh-like network around the nucleus, while vimentin filaments are predicted to appear more as fibrous filaments close to the cell boundary. (B) These predictions are consistent with experimental results where vimentin filaments form a mesh-like network cage around the nucleus, while their organization changes from mesh-like networks to fibrous structures with increasing distance from the cell center (*22*). Comparing simulations and experiments show that these fibrous vimentin filaments are aligned in the direction of the maximum tension in the cytoskeleton (scale bars: 10 μm and 5 μm in the top and bottom panels, respectively)

To test whether the state of mechanical stresses determined by the model impacts the spatial distribution of vimentin filaments, we compare the model predictions with the spatial organization of the vimentin network in living fibroblasts. It has been previously shown that “transcription activator-like effector nuclease” (TALEN)-based genome modification (*59*) can express fluorescent-labeled VIF in non-immortalized fibroblasts while minimizing changes induced in the VIF network due to tagging and overexpression of vimentin (*22*). In this TALEN-based genome editing method, the mEmerald gene is introduced at the N terminus of the endogenous vimentin locus to express fluorescent-labeled vimentin in fibroblasts. The spatial organization of the vimentin network can be then monitored by imaging the mEmerald-vimentin network at high resolution, and it has been shown that wild-type and TALEN-edited cells exhibit similar vimentin organization and cellular morphology with no detectable differences (*22*). These experiments show that, consistent with the high compression level around the nucleus predicted by our simulations, vimentin appears as wavy fibers and form a mesh-like network cage around the nucleus (Figure 5B, top panel). Also, it is shown that the mesh-like network structure gradually disappears with increasing distance from the nucleus and vimentin appears more as fibrous structures (Figure 5B, top panel). Together, these results validate the model prediction on the existence of two distinct families of intermediate filaments, namely, mesh-like and fibrous VIFs.

### Fibrous vimentin filaments are formed in the direction of the maximum principal stress in the cytoskeleton and extend more toward the cell periphery with increasing cytoskeletal tension

We next study the effect of the cytoskeletal stress field on the orientation of VIFs. To this end, we simulate contraction of a cell on a deformable substrate. As expected, our simulations show that the orientation of the substrate deformation vector *u* (black lines in Figure 5A) and traction force vector coincides with the direction of the maximum tensile stress vector *σ*^(t)^ (green lines in Figure 5A). On the other hand, TFM experiments of TALEN-edited cells depict that the orientation of traction forces on the substrate is highly aligned with that of local fibrous VIFs (Figure 5B, bottom panel) (*22*). Comparing our simulation with the TFM experiments show that fibrous VIFs are oriented in the direction of the stress vector *σ*^(t)^, indicating that this family of VIFs is aligned in the direction of the maximum tension in the cytoskeleton.

We next tested whether disruption of cytoskeletal tension affects the organization of fibrous VIFs. To this end, we culture cells on substrates with different stiffness (Figure S16) and we study how vimentin organization changes with reduction of cytoskeletal tension upon decreasing matrix stiffness. Simulating cells on soft and stiff substrates reveals that the maximum compressive stress *σ*^(c)^ remains high in the juxtanuclear region with softening of the substrate (purple line in Figure 6A), indicating the existence of mesh-like VIFs in the juxtanuclear region of cells on both soft and stiff substrates. The model prediction agrees with our experiments showing the appearance of wavy mesh-like VIFs in the juxtanuclear region which form a cage around the nucleus on both soft and stiff substrates (Figures 6B and S16). In contrast to the maximum compressive stress *σ*^(c)^, the model predicts that the maximum tensile stress *σ*^(t)^ significantly decreases in cells on the soft substrate (green line in Figure 6A). Concomitant with the reduction of tension in the fibrous VIFs predicted by the model, our experiments show that cells on the soft substrate have significantly lower numbers of fibrous VIFs and these filaments cannot reach the cell periphery (Figure 6B). Together, our results show that fibrous vimentin filaments are formed in the direction of the maximum principal stress in the cytoskeleton, and these fibrous filaments become more polymerized and extend toward the cell periphery with increasing cytoskeletal tension.

**Figure 6.**
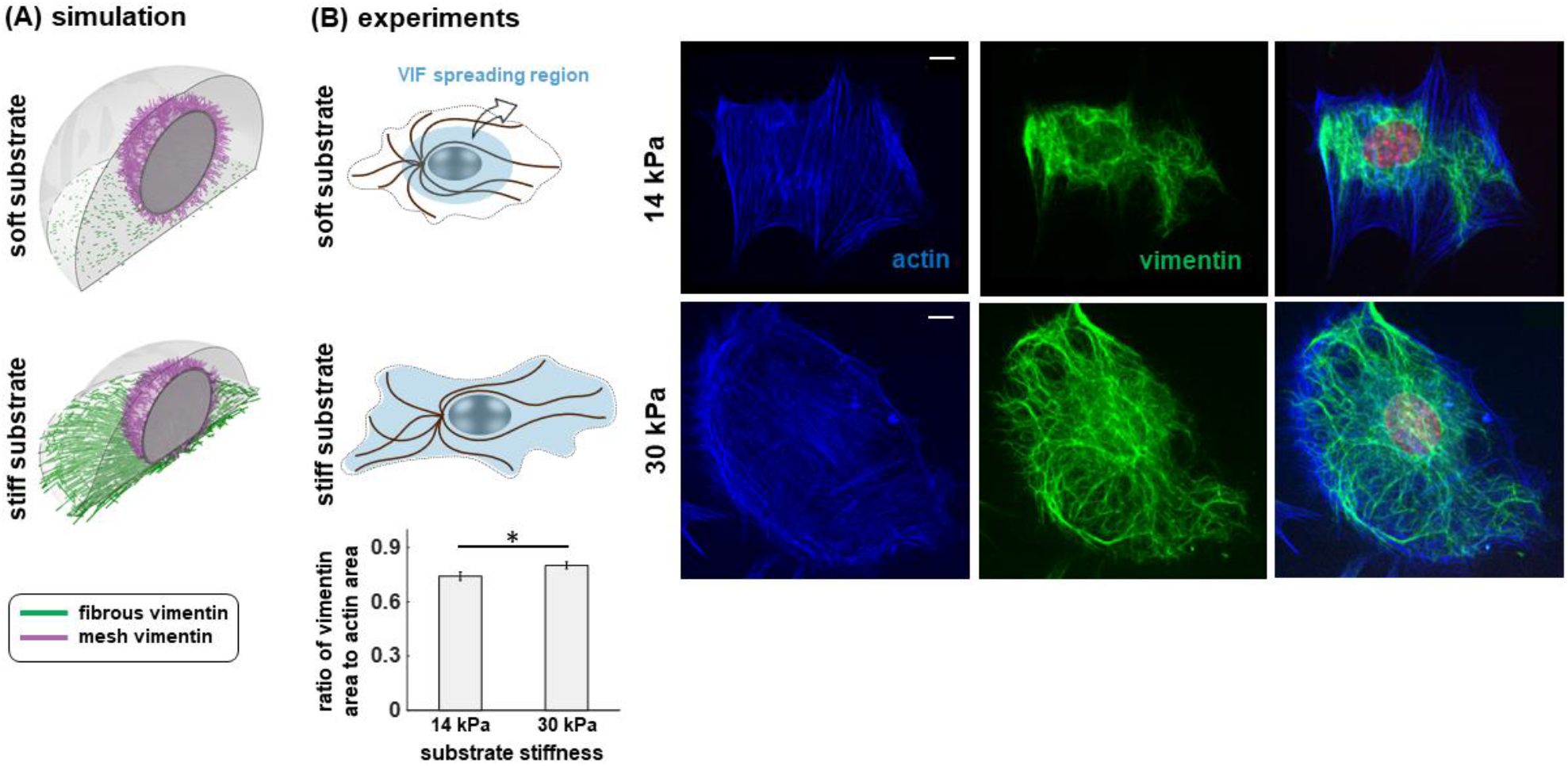
The effect of substrate stiffness on the formation of fibrous and mesh-like vimentin filaments. (A) To study how the vimentin filament organization changes with decreasing cytoskeletal tension, we simulate cells on soft and stiff circular micropatterned substrates. Our simulations show that, while the tensile stress *σ* ^(t)^ is remarkedly disrupted on soft substrates, cells on both substrates experience significant compressive stress *σ* ^(c)^ around the nucleus, expecting the existence of mesh-like VIFs in the juxtanuclear region of cells on both substrates. (B) In agreement with the simulations, our experiments show formation of mesh-like vimentin networks around the nucleus on both soft and stiff substrates. Furthermore, on soft substrates, concomitant with the reduction of tension in the fibrous VIFs predicted by the model, the vimentin network does not reach the cell periphery and cells lack fibrous vimentin filaments, indicating that fibrous VIFs become more polymerized and extend toward the cell periphery with increasing cytoskeletal tension. (*n* = 18-20, scale bars: 10 μm, error bars: standard error)

### Disruption of vimentin filaments affects microtubule organizations in a compression-dependent manner

We described that VIFs can laterally reinforce microtubules under actomyosin-based compressive stresses, and therefore disruption of VIFs can lead to higher compression and instability of microtubules in cells with high levels of actomyosin contractility (e.g., cells on stiff substrates) as shown in Figure 3D. To further test the effect of VIF depletion on microtubule organizations, we study the organization and density of microtubules in VIF +/+ and VIF −/− fibroblasts cultured on rigid micropatterned substrates. Our simulations show that, compared with VIF +/+ cells, microtubules in VIF −/− cells experience higher compression as they do not have the lateral reinforcement by the vimentin network against contractility-based compressive stresses (Figure 7A). Concomitant with the higher compression on microtubules predicted by the theoretical model, experimental results show that VIF −/− fibroblasts have lower microtubule densities compared with VIF +/+ cells (Figure 7B) (*60*). These results support the hypothesis that microtubules can be depolymerized under contractility-based compressive stresses when microtubules lose their lateral reinforcement upon vimentin depletion. Note that the symmetric geometry of cells in Figures 7A and 7B results in a symmetric distribution of compressive stresses on microtubules. Therefore, to ensure that vimentin depletion affects microtubule organizations in a compression-dependent manner, we study VIF +/+ and VIF −/− cells on an asymmetric micropatterned substrate where mechanical stresses are distributed asymmetrically in the cytoskeleton (Figures 7C-7E). Our simulations depict an asymmetric distribution of compressive stresses on microtubules when cells are cultured on a teardrop-like micropatterned geometry. These simulations illustrate that microtubules experience lower compression in region (ii) compared with region (i) (Figure 7E). Concomitant with the lower compression on microtubules in region (ii), experimental results exhibit higher microtubule densities in this region (*60*), consistent with our hypothesis that disrupting vimentin affects microtubule organizations in a compression-dependent manner. In conclusion, the 3D model enabled us to spatially map the stress field in the cytoskeleton of micropatterned cells which, in turn, identified new mechanisms through which vimentin coordinates microtubule organization.

**Figure 7.**
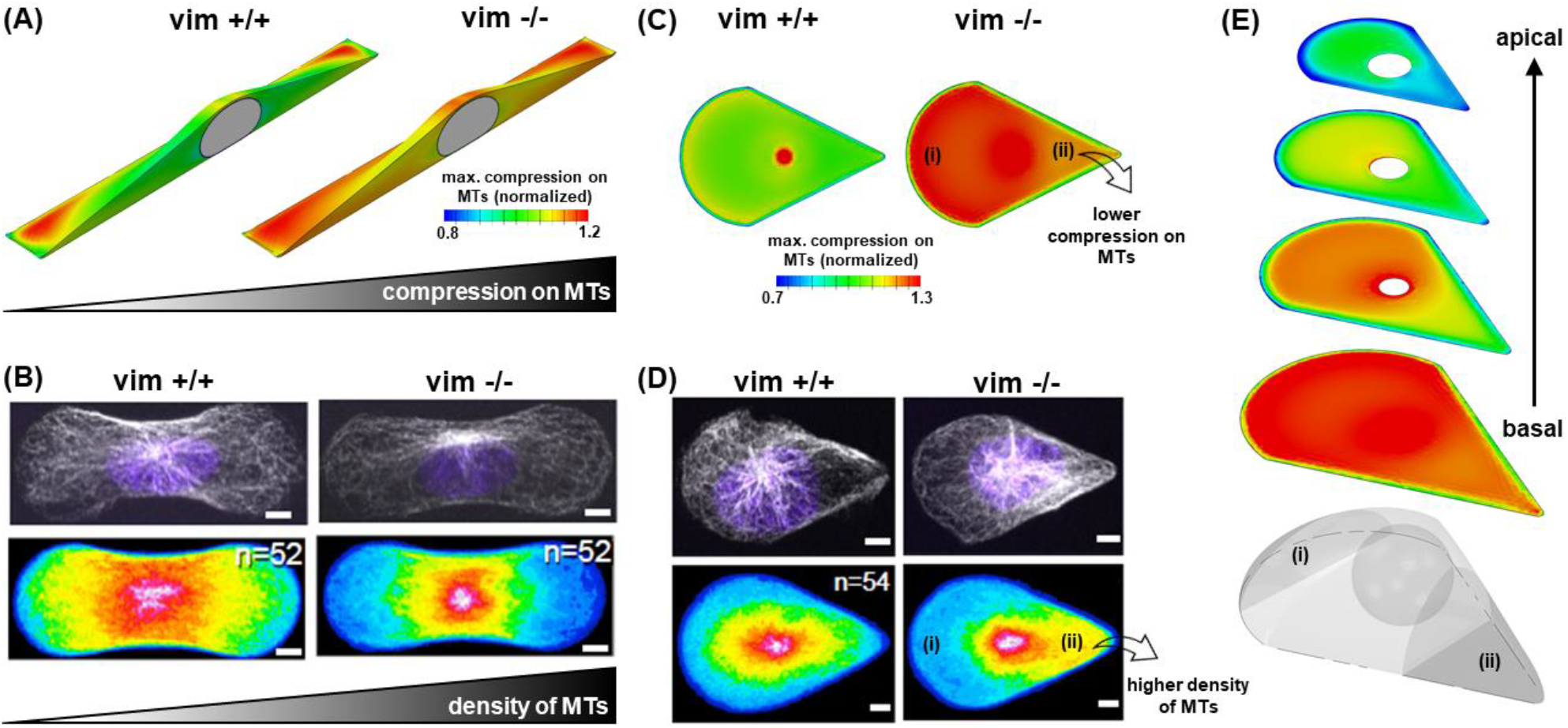
Disruption of vimentin filaments affects microtubule organizations in a compression-dependent manner. (A) Simulations of fibroblasts on rigid micropatterned substrates show that microtubules in VIF−/− cells experience higher compression than in control cells. (B) Concomitant with the higher compression on microtubules in VIF−/− cells, experimental results show lower microtubule densities in VIF−/− cells (*60*). (C and E) We next simulate fibroblasts on rigid micropatterned substrates with a teardrop shape which, unlike the rectangular geometry, has an asymmetric geometry. Like the simulations of the rectangular geometry, disruption of vimentin generates higher compression on microtubules. However, unlike the rectangular geometry, the teardrop geometry generates an asymmetric stress field where microtubules experience lower compression in region ii. (D) Concomitant with the lower compression on microtubules in region ii, experimental results show higher microtubule densities in this region (*60*). (Scale bars: 5 μm)

## Discussion and Conclusions

Our results elucidate that vimentin can exhibit matrix stiffness-dependent effects on traction forces as vimentin, directly and indirectly, interact with other cytoskeletal components (i.e., the actomyosin and microtubule networks) whose state and organization change with matrix stiffness. For example, cells on stiff substrates develop a strong contractile actomyosin network to transmit tensile forces to the substrate, and therefore depletion of vimentin does not significantly decrease the ability of the cytoskeleton to transmit the forces to the substrates (Figure S17B). On the other hand, microtubules experience high compression, and they require reinforcement from the vimentin network to withstand the compression. Thus, depletion of vimentin filaments can cause instability of the microtubule network, thereby increasing contractility and traction forces (Figure S17B). This prediction agrees well with experimental studies in the literature for fibroblasts on stiff substrates where VIF −/− cells generate significantly higher forces compared with VIF +/+ cells (*20–22*). In contrast, microtubules in cells on soft substrates experience low compression, and therefore depletion of vimentin filaments does not cause significant instability in the microtubule network (Figure S17A). On the other hand, contractility-associated vimentin filaments play an important role in the transmission of forces to the substrate as cells on soft substrates exhibit a weak actomyosin network. Therefore, depletion of vimentin in cells on soft substrates reduces traction forces (Figure S17A). This is consistent with other experimental results in the literature where knockout of vimentin in fibroblasts with low actomyosin contractility (e.g., cells on soft substrates) results in decreases in cell contractility and cell traction force (*17–19*).

The theoretical model accounts for myosin motors, actin filaments, microtubules, intermediate filaments, focal adhesions, and the nucleus which are the key cellular components involved in the generation and transmission of cellular forces to the ECM (Materials and Methods). In our coarse-grained model, myosin is represented by an active force-generating contractile element that is connected to the actin element in series, thereby generating tension in the actin element. Note that myosin contractile forces may also generate compression in actin filaments as observed in *in vitro* experimental models of actomyosin networks (*61*). However, the compressive stresses are relieved through buckling and severing of actin filaments, keeping only tensile f orces in the actin network (*61*). Therefore, we assume that the actin element only experiences tension in the model and transmits the tensile forces to the matrix through focal adhesions as observed experimentally (*34, 35, 62*).

The cell model responds to matrix stiffening by increasing myosin-generated contractile forces through a feedback mechanism between contractility and cytoskeletal tension (Figure S2). This agrees with experimental observations where the level of phosphorylated myosin motors in fibroblasts increases with matrix stiffness (*63*). Furthermore, concomitant with phosphorylation of more myosin motors and higher cell contractility, the actin element in the model stiffens in the direction of the tensile stresses, representing the recruitment and alignment of actin filaments in response to matrix stiffening (*7, 8, 33, 36*). Note that the formation of actin filaments in our simulations colocalizes with phosphorylation of myosin motors as observed experimentally (*32*). Starting with isotropic and uniform myosin and actin distributions, Figure S5 shows higher contractility and actin formation in basal regions and close to the cell boundary which is consistent with experimental observations (*32, 64, 65*). Disruption of the actomyosin network in the model, by inhibition of either myosin phosphorylation or actin formation, reduces cellular contractile forces as reported in experimental studies (*66*).

Microtubules in the model experience compression as the microtubule element is placed in parallel with the contractile myosin element. Although *in vitro* models show a complex interaction between the microtubules and actomyosin networks (*67*), the intrinsic cell contractility is known to generate compression in a large portion of microtubules which can in turn cause them to buckle (*12, 13*). Visualization of microtubule dynamics in cells transfected with GFP-tubulin shows that buckling of microtubules increases when cell contractility is stimulated by addition of thrombin to cells, while microtubules buckle less or even completely straighten with decreasing contractility after addition of cytochalasin D to destabilize the actin network (*11*). The placement of the microtubules in parallel with the active element is consistent with these experimental observations.

The model predicts that cell contractility, traction force, and matrix deformation increase with disruption of microtubules, which are all consistent with previous experimental studies (*41*). Kolodney and Elson showed that depolymerization of microtubules upon nocodazole treatment increases cell contractility by promoting phosphorylation of myosin light chains (*14*). The increase in cell contractility was found to be associated with stimulation of actin and focal adhesion organizations and formation of stress fibers (*42*). It was later shown that GEF-H1 is involved in this nocodazole-induced increase in contractility. GEF-H1 is a RhoA-specific guanine nucleotide exchange factor which is activated by microtubule depolymerization (*16*). The activated GEF-H1 activates the Rho-Rock pathway, which in turn increases phosphorylation of myosin motors and cell contractility. Consistent with the increase in cell contractility, other studie s showed that depolymerization of microtubules also leads to higher cell force generation in fibroblasts (*15, 43*) which generates larger strains in the underlying substrate (*68*).

The model contains two vimentin elements. The first element interacts with microtubules and stabilizes them under contractility-based compressive forces. Consistently, in vitro studies of isolated microtubules showed that microtubules without the VIF reinforcement are significantly less stable, buckle at much larger wave-lengths, and withstand remarkedly smaller compressive stresses compared with VIF-reinforced microtubules in living cells (*13*). Other experimental studies also showed that VIFs template and stabilize microtubule organizations as vimentin turns over much slower than microtubules (*46*). Furthermore, recent evidence from in vitro studies reveals that VIFs stabilize microtubules against depolymerization through direct physical interactions (*48*). All these studies show that vimentin filaments support and stabilize microtubules under contractility-based compressive stresses and prevent them from destabilization.

We discussed that in cells with high levels of actomyosin contractility, microtubules experience high actomyosin-based compression in the absence of vimentin filaments leading to instability (Figure 3D), reorganization (Figure 7D), and even depolymerization (Figure 7B) of microtubules. This agrees with recent studies on fibroblasts within 3D collagen matrices where disruption of actomyosin-based compressive forces on microtubules allows them to grow in length and number (*69*). In addition to fibroblasts, the same behavior has been recently reported in glioblastoma cells where disruption of actomyosin contractility, and subsequently compressive forces on microtubules, increase the length and number of microtubules (*70*). Our results are also supported by the studies on isolated microtubules where compression on microtubules reduces the rate of microtubule growth (*71, 72*) and increases the occurrence of microtubule catastrophe (*73*). Furthermore, depletion of VIFs in fibroblasts on stiff substrates has been reported to increase actomyosin contractility through activation of RhoA by GEF-H1 (*20*). As previously discussed, depolymerization of microtubules is known to activate GEF-H1 (*16*) which in turn activates the Rho-Rock pathway to increase phosphorylation of myosin motors and cell contractility (*14*). This may indicate that the increase in contractility upon VIF depletion can be due to depolymerization of microtubules. Note that immunoblot analyses of cell lysates from VIF +/+ and VIF −/− fibroblasts did not show a significant difference in microtubule expression levels (*74*). This indicates that the total amount of microtubules (both polymerized and depolymerized) measured by immunoblotting remains the same and the lack of vimentin in VIF −/− fibroblasts only affects microtubule stability, organization, and depolymerization rate as observed in Figure 3D, Figure 7D, and Figure 7B, respectively.

In addition to reinforcing microtubules under compression, vimentin is also known to interact with actin filaments. This interaction is captured by the second vimentin element in the model, and we show that the vimentin-actin interaction becomes more important in cells with low actomyosin levels (e.g., cells on soft substrates) as depletion of vimentin in these cells decreases contractile forces. This agrees with recent experimental studies showing that vimentin and actin filaments form an interpenetrating network, and lack of vimentin in VIF −/− fibroblasts on soft substrates reduces traction forces (*19*). This vimentin element in our model experiences tension which is consistent with experimental observations. Recently, in situ nonlinear Raman imaging of cells on 2D rigid substrates showed that vimentin filaments can be subject to tensile stresses due to the intrinsic contractility-driven cytoskeletal tension (*49*) which can, in turn, lead to unfolding of coiled-coil α-helical structures in vimentin filaments into anti-parallel β-strand structures (*75*). Furthermore, the tension-driven unfolding of vimentin filaments was found to decrease with disruption of cytoskeletal tension upon culturing cells on soft substrates or using actomyosin inhibitors including blebbistatin and latrunculin A (*49*). Similarly, vimentin filaments in “tensegrity models” are assumed to undergo tensile stresses (*11*). Also, “actomyosin-associated vimentin intermediate filaments” have been shown to play a critical role in the transmission of tensile stresses between the nucleus and the ECM (*76*). All these studies show that vimentin filaments are involved in the transmission of contractility-based tensile forces to the ECM through interactions with contractile actin filaments, and our results show that the balance between vimentin-actin and vimentin-microtubule interactions regulates the effect of vimentin on cellular forces.

Taken together, our study elucidates the complex crosstalk between vimentin, actomyosin, and microtubules which impacts cell-generated traction forces in a matrix stiffness-dependent manner. Vimentin is involved in various important biological processes including migration (*77*), polarity (*78*), EMTs (*79*), cataracts (*80*), and cancer progression (*81*). Given that cellular traction forces are central to both wound healing and a wide range of pathological processes including fibrosis and surgical adhesions, our study has broad implications for understanding the effect of vimentin on cell-generated traction forces within different physiological and pathological microenvironments.

## ACKNOWLEDGMENTS

We acknowledge helpful discussions with Gaudenz Danuser. This work was supported by National Cancer Institute Award R01CA232256 and U54CA261694; National Institute of Biomedical Imaging and Bioengineering Awards R01EB017753 and R01EB030876; National Institute of General Medicine award R01GM096971, the NSF Center for Engineering Mechanobiology Grant CMMI-154857; NSF Grants MRSEC/DMR-1720530 and DMS-1953572.

## Author contributions

F. Alisafaei and V. B. Shenoy designed the research. K. Mandal, M. Swoger, H. Yang, M. Guo, P. A. Janmey, and A. E. Patteson performed and supervised the experiments. F. Alisafaei and V. B. Shenoy developed the theoretical models and performed numerical simulations. F. Alisafaei, and V. B. Shenoy wrote the manuscript and all authors discussed the results and commented on the manuscript.

## Materials and Methods

### Cell culture, reagents, immunostaining

Wild type or vimentin null mouse embryonic fibroblast (MEF) cells are grown in 1X DMEM (Life Technologies) supplemented with 10% (vol/vol) FBS (GE Healthcare Life Sciences), 1% nonessential amino acid and 10mM HEPES at 37 °C with 5% (vol/vol) CO2. Cells are plated at a density of 10,000 cells/gel (18mm coverslip) or less.

For immunofluorescence experiments, cells are fixed with 4% paraformaldehyde (Affymetrix) followed by 5% BSA and 1% Saponin (Sigma) for blocking and permeabilization. Primary antibodies are Alexa-Fluor 647 phalloidin (Invitrogen), and anti-vimentin (Novus Biologicals), dapi (Sigma).

### Microscopy and Imaging

Images of the cell and beads are acquired with a Leica DMIRE2 microscope using iVision software. An environmental chamber is used to maintain the temperature at 37 °C and 5% CO2 for live-cell imaging. Bright-field images of cells and fluorescent images of the beads are acquired at multiple positions with a 40X objective. Cells were imaged by manual tracking using Fiji at a rate of 10 mins per frame.

### Cell traction force microscopy

To perform traction force microscopy experiments, polyacrylamide hydrogel substrates of desired stiffness with 1% of 200 nm fluorescently labeled green beads (2% solid, Thermo Fisher Scientific) were prepared as described previously (*82*). The concentration of acrylamide and bisacrylamide varied for different matrix stiffness. Substrates are coated with 50 ug/ml of either collagen type I (Corning) or fish fibronectin (homemade). After 24 hrs of plating cells, phase images of the cell, stressed and relaxed images of fluorescently labeled beads were acquired using an epifluorescence microscope with an environmental chamber. For the traction force microscopy analysis, a custom-built Matlab code was used. At any time, the exerted force can be calculated from the bead displacement field using constrained Fourier Transform Traction Microscopy (*83*). The details of the calculation can be found in (*84*). Image acquisition is performed as we maintain 37°C and 5% CO_2_ throughout the experiments.

### Theoretical model

We developed a theoretical cell model for three-dimensional and one-dimensional frameworks as described in detail in SI Sections 1 and 2, respectively. The cell model includes the following components: the cytoskeleton, the focal adhesions, and the nucleus. As described in the main text, the cytoskeleton is composed of the myosin motors, the microtubules, the actin filaments, and the vimentin filaments (see SI for more details and Table S1 for the model parameters used in our simulations). Focal adhesions in our coarse-grained model are treated as a set of initially soft nonlinear mechanical elements that stiffen with tension to capture the tension-dependent formation of the focal adhesions. When the tensile stress exerted by the contractile cell to the adhesion layer exceeds a certain threshold, mature focal adhesions are formed, and the cell is connected to the substrate, while below this threshold, the stiffness of the adhesion layer remains low and the substrate experiences negligible forces. The nucleus is modeled as a fibrous elastic thin layer (representing the nuclear envelope and lamina layer) filled with a solid material which represents chromatin and other subnuclear components (see (*32*) for more details). The matrix is modeled as a thick linear elastic material. The theoretical model focuses on the stationary behavior of fibroblasts where cells have fully spread on the substrate. Note that the model can be readily extended to study the time-dependent behavior of cells as described in our recent publications (*85*). Similarly, our experiments are performed in the stationary configuration where cells have fully spread on the substrate.

## Supplementary Information

### Three-dimensional cell model

Each component of the model is described in detail and the model is implemented into a three-dimensional finite element framework.

### One-dimensional cell model

The model presents the key features of the three-dimensional model without the complexity of the finite element implementation.

**Supplementary references S1-S21**

**Figures S1-S18**

**Table S1:** List of parameters used in the model.

