## Supplementary Information for "Vimentin Intermediate Filaments Can Enhance or Abate Active Cellular Forces in a Microenvironmental Stiffness-Dependent Manner"

[1] Department of Mechanical and Industrial Engineering, New Jersey Institute of Technology, Newark, NJ 07102 USA
[2] Center for Engineering Mechanobiology, University of Pennsylvania, Philadelphia, PA 19104 USA
[3] Institute for Medicine and Engineering, University of Pennsylvania, 3340 Smith Walk, Philadelphia, PA 19104 USA
[4] Physics Department, Syracuse University, Syracuse, New York 13244 USA
[5] BioInspired Institute, Syracuse University, Syracuse, New York 13244 USA
[6] Department of Mechanical Engineering, Massachusetts Institute of Technology, Cambridge, MA 02139 USA
[7] Departments of Physiology, and Physics & Astronomy, University of Pennsylvania, Philadelphia, PA 19104 USA
[8] Department of Materials Science and Engineering, School of Engineering and Applied Science, University of Pennsylvania, Philadelphia, PA 19104 USA


### Supplementary Information

#### 1. Three-dimensional cell model

The model is composed of the following elements: (i) the myosin molecular motors, (ii) the microtubules, (iii) the actin filaments, and (iv) the vimentin filaments. We first describe the model without the presence of the vimentin element and we will later describe how the vimentin filament network can be added to the model.

##### 1.1. Myosin motors, microtubules, and actin filaments

Here, we first define a contractility tensor, $\rho_{ij}$, whose components represent cell contractility in different directions. We then discuss the properties of the contractility tensor and the mechanisms through which $\rho_{ij}$ changes with the mechanical properties of the extracellular matrix. At submicron levels, the force generated by a myosin motor can be treated as a force dipole (Figure 1C) which is a pair of equal but oppositely directed forces $F_i(x_j)$ and $-F_i(x_j+\Delta x_j)$ where $x_j$ and $x_j+\Delta x_j$ are coordinates of myosin head domains, and $|\Delta x_j|$ is the length of myosin II filaments (approximately 200 nm) (*1*). The work done by the force dipole can be determined as follows

$$W_{\text{dipole}} = F_i u_i(x_j+\Delta x_j) - F_i u_i(x_j) \tag{S1.1}$$

where $u_i(x_j)$ and $u_i(x_j+\Delta x_j)$ are the displacements of the cytoskeleton at $x_j$ and $x_j+\Delta x_j$, respectively. We then can calculate the total work generated by all myosin motors per volume $V$

$$W = \left(\frac{1}{V}\right)\sum_{k=1}^{N}\left[F_i^{(k)} u_i\left(x_j^{(k)} + \Delta x_j^{(k)}\right) - F_i^{(k)} u_i\left(x_j^{(k)}\right)\right] \tag{S1.2}$$

and rewrite it in this form

$$W = \left(\frac{1}{V}\right)\sum_{k=1}^{N} F_i^{(k)}\,\Delta x_j^{(k)}\,\partial_j u_i \tag{S1.3}$$

using the following definition

$$\partial_j u_i = \frac{u_i\left(x_j^{(k)} + \Delta x_j^{(k)}\right) - u_i\left(x_j^{(k)}\right)}{\Delta x_j^{(k)}} \tag{S1.4}$$

where $N$ is the total number of phosphorylated (bound) myosin motors. Finally, we can write the total work as follows

$$W = \rho_{ij}\varepsilon_{ij} \tag{S1.5}$$

where

$$\varepsilon_{ij} = \frac{1}{2}\left(\partial_j u_i + \partial_i u_j\right) \tag{S1.6}$$

is the linearized strain, and

$$\rho_{ij} = \left(\frac{1}{V}\right)\sum_{k=1}^{N} F_i^{(k)}\,\Delta x_j^{(k)} \tag{S1.7}$$

represents the cell contractility which is a symmetric tensor as the following relationship holds

$$F_i^{(k)}\,\Delta x_j^{(k)} = F_j^{(k)}\,\Delta x_i^{(k)} \tag{S1.8}$$

Equation (S1.7) shows that $\rho_{ij}$ is related to the density of phosphorylated myosin molecular motors. In our coarse-grain model, we therefore use $\rho_{ij}$ as a tensor whose components represent cell contractility in different directions. Experimental studies show that the magnitude and the direction of cellular contractility depend on the physical properties of the microenvironment. For example, cells cultured on stiff microenvironments are more contractile and have higher densities of phosphorylated myosin than those cultured on soft microenvironments (*2*). As discussed in the main text, to define the cell contractility tensor $\rho_{ij}$ in our coarse-grain model, we hypothesize that the average of contractility in all three directions, $\frac{1}{3}\rho_{kk} = (\rho_{11} + \rho_{22} + \rho_{33})/3$, increases with the average of tension in the actin filament network, $\frac{1}{3}\sigma_{kk} = (\sigma_{11} + \sigma_{22} + \sigma_{33})/3$

$$\frac{\rho_{kk}}{3} = f_{\mathrm{m}}\frac{\sigma_{kk}}{3} + f_0\,\rho_0 \tag{S1.9}$$

where this stress-dependent feedback mechanism is regulated by the feedback parameter $f_{\mathrm{m}}$, $\rho_0$ is the initial contractility (basal cell contractility), and $f_0$ regulates the mean contractility $\frac{1}{3}\rho_{kk}$ in the absence of tension ($\sigma_{kk} = 0$).

To implement the model presented in (S1.9) into a three-dimensional finite element framework, we need to define the stress tensor $\sigma_{ij}$ and the stiffness tensor $\mathrm{C}_{ij}$. In what follows, we first derive these equations and we then show that as a result of the feedback mechanism in equation (S1.9), in addition to the cell contractility $\rho_{ij}$, the stiffness of the actin network $\mathrm{C}_{ijkl}^{(\mathrm{A})}$ and the tension it carries, $\sigma_{ij}$, also increase with matrix stiffness in an orientation-dependent manner. To this end, we start with the following definition for $\rho_{ij}$ which relates it to the strain tensor $\boldsymbol{\varepsilon}^{(\mathrm{X})}$ (the three-dimensional representation of $\varepsilon^{(\mathrm{X})}$ shown in Figure S18)

$$\rho_{ij} = K^{(\rho)}\varepsilon_{kk}^{(\mathrm{X})}\delta_{ij} + 2\mu^{(\rho)}\left(\varepsilon_{ij}^{(\mathrm{X})} - \frac{1}{3}\varepsilon_{kk}^{(\mathrm{X})}\delta_{ij}\right) + \bar{\rho}_0\delta_{ij} \tag{S1.10}$$

where

$$K^{(\rho)} = \frac{3K^{(\mathrm{MT})}\alpha_{\mathrm{v}} - 1}{3(\beta_{\mathrm{v}} - \alpha_{\mathrm{v}})} \tag{S1.11}$$

is the motor density effective modulus,

$$\mu^{(\rho)} = \frac{2\mu^{(\mathrm{MT})}\alpha_{\mathrm{d}} - 1}{2(\beta_{\mathrm{d}} - \alpha_{\mathrm{d}})} \tag{S1.12}$$

is the polarization effective modulus,

$$\bar{\rho}_0 = \frac{\beta_{\mathrm{v}}\,\rho_0}{\beta_{\mathrm{v}} - \alpha_{\mathrm{v}}} \tag{S1.13}$$

is the effective contractility,

$$K^{(\mathrm{MT})} = \frac{E^{(\mathrm{MT})}}{3(1 - 2v^{(\mathrm{MT})})} \tag{S1.14}$$

is the bulk modulus of the cytoskeletal components that are in compression (*e.g.*, microtubule network), and

$$\mu^{(\mathrm{MT})} = \frac{E^{(\mathrm{MT})}}{2(1 + v^{(\mathrm{MT})})} \tag{S1.15}$$

is the shear modulus of the cytoskeletal components that are in compression (*e.g.*, microtubule network). In the above equations, $E^{(\mathrm{MT})}$ is the elastic modulus of the cytoskeletal components that are in compression (*e.g.*, microtubule network), $v^{(\mathrm{MT})}$ is the Poisson's ratio of the cytoskeletal components that are in compression (*e.g.*, microtubule network), $\rho_0$ is the initial contractility, $\alpha_{\mathrm{v}}$ is the volumetric chemo-mechanical feedback parameter (large values of $\alpha_{\mathrm{v}}$ lead to higher densities of phosphorylated myosin motors), $\alpha_{\mathrm{d}}$ is the deviatoric chemo-mechanical feedback parameter which represents the tendency of the cell to generate polarized contraction (small values of $\alpha_{\mathrm{d}}$ lead to non-polarized contractility), $\beta_{\mathrm{v}}$ is the volumetric chemical stiffness parameter which regulates the mean contractility $\frac{1}{3}\rho_{kk}$ in the absence of tension ($\sigma_{kk} = 0$) by maintaining the contractility at a basal level (large values of $\beta_{\mathrm{v}}$ make phosphorylation of myosin more difficult), $\beta_{\mathrm{d}}$ is the deviatoric chemical stiffness parameter which represents the disinclination of myosin to orient along the cell's polarization direction (large values of $\beta_{\mathrm{d}}$ cause myosin motors to orient randomly).

Consistent with experimental observations in references (*3*, *4*) which show that microtubules are compressed by the internally generated cell contractile forces, the cell contractility $\rho_{ij}$ generates the compressive stress $\mathrm{C}_{ijkl}^{(\mathrm{MT})}\varepsilon_{kl}^{(\mathrm{X})}$ on the microtubule network where

$$\mathrm{C}_{ijkl}^{(\mathrm{MT})} = K^{(\mathrm{MT})}\delta_{ij}\delta_{kl} + \mu^{(\mathrm{MT})}\left(\delta_{ik}\delta_{jk} + \delta_{il}\delta_{jk} - \frac{2}{3}\delta_{ij}\delta_{kl}\right) \tag{S1.16}$$

is the stiffness tensor of the microtubule network. In addition to compressively loading the microtubule network, the cell contractility $\rho_{ij}$ also generates tensile stress in the actin filament network, $\sigma_{ij}$,

$$\rho_{ij} = -\mathrm{C}_{ijkl}^{(\mathrm{MT})}\varepsilon_{kl}^{(\mathrm{X})} + \sigma_{ij} \tag{S1.17}$$

Equation (S1.10) indicates that the contractility tensor $\rho_{ij}$ is initially isotropic and, as a result, the cell exhibits the same contractility in all directions in the initial configuration. This can be mathematically shown by rewriting equation (S1.10) in the following form

$$\rho_{ij} = \mathrm{C}_{ijkl}^{(\rho)}\varepsilon_{kl}^{(\mathrm{X})} + \bar{\rho}_0\delta_{ij} \tag{S1.18}$$

where

$$\mathrm{C}_{ijkl}^{(\rho)} = K^{(\rho)}\delta_{ij}\delta_{kl} + \mu^{(\rho)}\left(\delta_{ik}\delta_{jk} + \delta_{il}\delta_{jk} - \frac{2}{3}\delta_{ij}\delta_{kl}\right) \tag{S1.19}$$

Equations (S1.17) and (S1.18) show that, in the stress-free state $\sigma_{ij} = 0$ (initially tension is zero), the diagonal components of the contractility tensor $\rho_{ij}$ are all equal and non-zero $\rho_{11} = \rho_{22} =$

$\rho_{33} \neq 0$, while the off-diagonal components are all zero $\rho_{12} = \rho_{21} = \rho_{13} = \rho_{31} = \rho_{23} = \rho_{32} = 0$ indicating that $\rho_{ij}$ is initially isotropic. This can be better shown by writing equation (S1.18) in Voigt notation

$$\begin{Bmatrix} \rho_{11} \\ \rho_{22} \\ \rho_{33} \\ \rho_{12} \\ \rho_{13} \\ \rho_{23} \end{Bmatrix} = \begin{bmatrix} C^{(\rho)}_{1111} & C^{(\rho)}_{1122} & C^{(\rho)}_{1133} & C^{(\rho)}_{1112} & C^{(\rho)}_{1113} & C^{(\rho)}_{1123} \\ C^{(\rho)}_{2211} & C^{(\rho)}_{2222} & C^{(\rho)}_{2233} & C^{(\rho)}_{2212} & C^{(\rho)}_{2213} & C^{(\rho)}_{2223} \\ C^{(\rho)}_{3311} & C^{(\rho)}_{3322} & C^{(\rho)}_{3333} & C^{(\rho)}_{3312} & C^{(\rho)}_{3313} & C^{(\rho)}_{3323} \\ C^{(\rho)}_{1211} & C^{(\rho)}_{1222} & C^{(\rho)}_{1233} & C^{(\rho)}_{1212} & C^{(\rho)}_{1213} & C^{(\rho)}_{1223} \\ C^{(\rho)}_{1311} & C^{(\rho)}_{1322} & C^{(\rho)}_{1333} & C^{(\rho)}_{1312} & C^{(\rho)}_{1313} & C^{(\rho)}_{1323} \\ C^{(\rho)}_{2311} & C^{(\rho)}_{2322} & C^{(\rho)}_{2333} & C^{(\rho)}_{2312} & C^{(\rho)}_{2313} & C^{(\rho)}_{2323} \end{bmatrix} \begin{Bmatrix} \varepsilon_{11} \\ \varepsilon_{22} \\ \varepsilon_{33} \\ \varepsilon_{12} \\ \varepsilon_{13} \\ \varepsilon_{23} \end{Bmatrix} + \begin{Bmatrix} \bar{\rho}_0 \\ \bar{\rho}_0 \\ \bar{\rho}_0 \\ 0 \\ 0 \\ 0 \end{Bmatrix} \tag{S1.20}$$

where the first, second, and third components of the last term in equation (S1.20) have the same value $\bar{\rho}_0$.

In addition to the feedback mechanism between contractility and tension in equation (S1.9) and consistent with experimental observations (*5*), we also hypothesize that the stiffness of the actin network $C^{(A)}_{ijkl}$ increases in proportion and in the directions of the tensile principal components of the stress tensor $\sigma_{ij}$, (*6*)

$$C^{(A)}_{ijkl} = C^{(I)}_{ijkl} + C^{(F)}_{ijkl} \tag{S1.21}$$

where $\mathbf{C}^{(I)}$ is the initial stiffness of the actin filaments network and $\mathbf{C}^{(F)}$ denotes the stiffening of the actin network with tension (but not in compression). The initial stiffness of the actin network $\mathbf{C}^{(I)}$ is defined as follows

$$C^{(I)}_{ijkl} = K^{(I)}\delta_{ij}\delta_{kl} + \mu^{(I)}\left(\delta_{ik}\delta_{jk} + \delta_{il}\delta_{jk} - \frac{2}{3}\delta_{ij}\delta_{kl}\right) \tag{S1.22}$$

where

$$K^{(I)} = \frac{E^{(I)}}{3(1-2v^{(I)})} \tag{S1.23}$$

is the initial bulk modulus of the actin network, and

$$\mu^{(I)} = \frac{E^{(I)}}{2(1+v^{(I)})} \tag{S1.24}$$

is the initial shear modulus of the actin network, $E^{(I)}$ is the initial elastic modulus of the actin network, and $v^{(I)}$ is the initial Poisson's ratio of the actin network.

Next, we define the stiffening part of the stiffness tensor, $\mathbf{C}^{(F)}$. To this end, we first decompose $\boldsymbol{\sigma}$

$$\sigma_{ij} = \sigma^{(A)}_{ij} = \sigma^{(I)}_{ij} + \sigma^{(F)}_{ij} \tag{S1.25}$$

where $\boldsymbol{\sigma}^{(I)}$

$$\sigma^{(I)}_{ij} = C^{(I)}_{ijkl}\,\varepsilon^{(Y)}_{kl} \tag{S1.26}$$

is linearly related to the strain tensor $\boldsymbol{\varepsilon}^{(Y)}$ (the three-dimensional representation of $\varepsilon^{(Y)}$ shown in Figure S18) which can be written as a function of its eigenvalues (principal strains) $\varepsilon^{(Y)}_1$, $\varepsilon^{(Y)}_2$, $\varepsilon^{(Y)}_3$ and eigenvectors $\boldsymbol{n}_1$, $\boldsymbol{n}_2$, $\boldsymbol{n}_3$

$$\boldsymbol{\varepsilon}^{(Y)} = \sum_{i=1}^{3} \varepsilon^{(Y)}_i \boldsymbol{n}_i \otimes \boldsymbol{n}_i = \sum_{i=1}^{3} \varepsilon^{(Y)}_i \mathbf{E}_i \tag{S1.27}$$

where the symmetric tensors $\mathbf{E}_1 = \boldsymbol{n}_1 \otimes \boldsymbol{n}_1$, $\mathbf{E}_2 = \boldsymbol{n}_2 \otimes \boldsymbol{n}_2$, and $\mathbf{E}_3 = \boldsymbol{n}_3 \otimes \boldsymbol{n}_3$ are the eigenprojections of $\boldsymbol{\varepsilon}^{(Y)}$ and $\otimes$ denotes the dyadic product of two arbitrary vectors $\boldsymbol{u}$ and $\boldsymbol{v}$ as $(\boldsymbol{u} \otimes \boldsymbol{v})_{ij} = u_i v_j$. With the eigenvalues ($\varepsilon^{(Y)}_1$, $\varepsilon^{(Y)}_2$, $\varepsilon^{(Y)}_3$) and eigenvectors ($\boldsymbol{n}_1$, $\boldsymbol{n}_2$, $\boldsymbol{n}_3$) at hand, we next define $\sigma^{(F)}_{ij}$ in (S1.25)

$$\boldsymbol{\sigma}^{(\mathrm{F})} = \sum_{i=1}^{3} \frac{\partial f(\varepsilon_i^{(\mathrm{Y})})}{\partial \varepsilon_i^{(\mathrm{Y})}} \boldsymbol{n}_i \otimes \boldsymbol{n}_i = \sum_{i=1}^{3} \sigma^{(\mathrm{F})}\left(\varepsilon_i^{(\mathrm{Y})}\right) \mathbf{E}_i = \sum_{i=1}^{3} \sigma_i^{(\mathrm{F})} \mathbf{E}_i \tag{S1.28}$$

where $\sigma_i^{(\mathrm{F})}$ are the eigenvalues (principal stresses) of the stress tensor $\sigma_{ij}^{(\mathrm{F})}$ and are defined as follows

$$\sigma_i^{(\mathrm{F})} = \frac{\partial f(\varepsilon_i^{(\mathrm{Y})})}{\partial \varepsilon_i^{(\mathrm{Y})}} = \tag{S1.29}$$

$$\begin{cases} 0 & \varepsilon_i^{(\mathrm{Y})} < \epsilon_1 \\ \ell \dfrac{\left(\dfrac{\varepsilon_i^{(\mathrm{Y})} - \epsilon_1}{\epsilon_2 - \epsilon_1}\right)^t \left(\varepsilon_i^{(\mathrm{Y})} - \epsilon_1\right)^2}{(t+1)(t+2)} & \epsilon_1 \le \varepsilon_i^{(\mathrm{Y})} < \epsilon_2 \\ \ell \left[ \dfrac{\left(1 + \varepsilon_i^{(\mathrm{Y})} - \epsilon_2\right)^{s+2} - 1}{(s+1)(s+2)} + \dfrac{\epsilon_2 - \varepsilon_i^{(\mathrm{Y})}}{s+1} + \dfrac{(\varepsilon_i^{(\mathrm{Y})} - \epsilon_2)(\epsilon_2 - \epsilon_1)}{t+1} + \dfrac{(\epsilon_2 - \epsilon_1)^2}{(t+1)(t+2)} \right] & \varepsilon_i^{(\mathrm{Y})} \ge \epsilon_2 \end{cases}$$

to ensure the continuity and smoothness of the first and second derivatives of $\sigma_i^{(\mathrm{F})}$ with respect to $\varepsilon_i^{(\mathrm{Y})}$ at the transition points $\epsilon_1 = \epsilon_\mathrm{c} - 0.5\epsilon_\mathrm{t}$ and $\epsilon_2 = \epsilon_\mathrm{c} + 0.5\epsilon_\mathrm{t}$ where $\epsilon_\mathrm{t} = 0.25\epsilon_\mathrm{c}$ is the transition width, $\epsilon_\mathrm{c}$ is the critical (tensile) principal strain, and $t$ is the transition constant. Equation (S1.29) shows that for large tensile strains $\varepsilon_i^{(\mathrm{Y})} \ge \epsilon_2$, the principal stress $\sigma_i^{(\mathrm{F})}$ nonlinearly increases with the principal strain $\varepsilon_i^{(\mathrm{Y})}$ where this increase is regulated by the stiffening parameters $\ell$ and $s$. With $\boldsymbol{\sigma}^{(\mathrm{F})}$ at hand from equation (S1.28), we can now determine $\mathbf{C}^{(\mathrm{F})}$ as follows

$$\mathrm{C}_{ijkl}^{(\mathrm{F})} = \frac{\mathrm{d}\sigma_{ij}^{(\mathrm{F})}}{\mathrm{d}\varepsilon_{kl}^{(\mathrm{Y})}} \qquad \text{or} \qquad \mathbf{C}^{(\mathrm{F})} = \frac{\mathrm{d}\boldsymbol{\sigma}^{(\mathrm{F})}}{\mathrm{d}\boldsymbol{\varepsilon}^{(\mathrm{Y})}} \tag{S1.30}$$

The piecewise linear approximation in equation (S1.30) requires $\sigma_{ij}^{(\mathrm{F})}$ as a function of $\varepsilon_{ij}^{(\mathrm{Y})}$ while equation (S1.29) gives $\sigma_i^{(\mathrm{F})}$ as a function of $\varepsilon_i^{(\mathrm{Y})}$. Therefore, we first write $\mathrm{C}_{ijkl}^{(\mathrm{F})}$ in the following form using the definition of $\sigma_{ij}^{(\mathrm{F})}$ in equation (S1.28)

$$\mathbf{C}^{(\mathrm{F})} = \sum_{i=1}^{3} \left\{ \mathbf{E}_i \otimes \frac{\mathrm{d}\sigma_i^{(\mathrm{F})}}{\mathrm{d}\boldsymbol{\varepsilon}^{(\mathrm{Y})}} + \sigma_i^{(\mathrm{F})} \frac{\mathrm{d}\mathbf{E}_i}{\mathrm{d}\boldsymbol{\varepsilon}^{(\mathrm{Y})}} \right\} \tag{S1.31}$$

and we then expand the first term in the right-hand side of equation (S1.31) by applying the chain rule

$$\mathbf{C}^{(\mathrm{F})} = \sum_{i=1}^{3} \left\{ \sum_{j=1}^{3} \frac{\partial \sigma_i^{(\mathrm{F})}}{\partial \varepsilon_j^{(\mathrm{Y})}} \mathbf{E}_i \otimes \frac{\mathrm{d}\varepsilon_j^{(\mathrm{Y})}}{\mathrm{d}\boldsymbol{\varepsilon}^{(\mathrm{Y})}} + \sigma_i^{(\mathrm{F})} \frac{\mathrm{d}\mathbf{E}_i}{\mathrm{d}\boldsymbol{\varepsilon}^{(\mathrm{Y})}} \right\} \tag{S1.32}$$

To further expand equation (1.32) and derive an analytical form for $\mathbf{C}^{(\mathrm{F})}$, we consider the three following cases. In the first case, the three eigenvalues of the strain tensor $\varepsilon_{ij}^{(\mathrm{Y})}$ are all nonidentical ($\varepsilon_1^{(\mathrm{Y})} \neq \varepsilon_2^{(\mathrm{Y})} \neq \varepsilon_3^{(\mathrm{Y})}$). In this case, we can derive the following analytical expression for $\mathbf{C}^{(\mathrm{F})}$ from equation (S1.32) by taking the derivatives of $\varepsilon_j^{(\mathrm{Y})}$ and $\mathbf{E}_i$ with respect to $\boldsymbol{\varepsilon}^{(\mathrm{Y})}$

$$\mathbf{C}^{(\mathrm{F})} = \sum_{a=1}^{3} \frac{\sigma_a^{(\mathrm{F})}}{(\varepsilon_a^{(\mathrm{Y})} - \varepsilon_b^{(\mathrm{Y})})(\varepsilon_a^{(\mathrm{Y})} - \varepsilon_c^{(\mathrm{Y})})} \left\{ \frac{\mathrm{d}\left(\boldsymbol{\varepsilon}^{(\mathrm{Y})}\right)^2}{\mathrm{d}\boldsymbol{\varepsilon}^{(\mathrm{Y})}} - \left(\varepsilon_b^{(\mathrm{Y})} + \varepsilon_c^{(\mathrm{Y})}\right)\mathbf{I}_\mathrm{S} - \left[\left(\varepsilon_a^{(\mathrm{Y})} - \varepsilon_b^{(\mathrm{Y})}\right) + \left(\varepsilon_a^{(\mathrm{Y})} - \varepsilon_c^{(\mathrm{Y})}\right)\right] \mathbf{E}_a \otimes \mathbf{E}_a - \left(\varepsilon_b^{(\mathrm{Y})} - \varepsilon_c^{(\mathrm{Y})}\right)(\mathbf{E}_b \otimes \mathbf{E}_b - \mathbf{E}_c \otimes \mathbf{E}_c) \right\} + \sum_{i=1}^{3}\sum_{j=1}^{3} \frac{\partial \sigma_i^{(\mathrm{F})}}{\partial \varepsilon_j^{(\mathrm{Y})}} \mathbf{E}_i \otimes \mathbf{E}_j \quad \text{(S1.33)}$$

where

$$\left(\frac{\mathrm{d}\left(\boldsymbol{\varepsilon}^{(\mathrm{Y})}\right)^2}{\mathrm{d}\boldsymbol{\varepsilon}^{(\mathrm{Y})}}\right)_{ijkl} = \frac{1}{2}\left(\delta_{ik}\varepsilon_{lj}^{(\mathrm{Y})} + \delta_{il}\varepsilon_{kj}^{(\mathrm{Y})} + \delta_{jl}\varepsilon_{ik}^{(\mathrm{Y})} + \delta_{kj}\varepsilon_{il}^{(\mathrm{Y})}\right) \quad \text{(S1.34)}$$

is the derivative of the square of $\boldsymbol{\varepsilon}^{(\mathrm{Y})}$, and

$$(\mathbf{I}_\mathrm{S})_{ijkl} = \frac{1}{2}\left(\delta_{ik}\delta_{jl} + \delta_{il}\delta_{jk}\right) \quad \text{(S1.35)}$$

is the symmetric identity tensor. In the second case, the strain tensor $\varepsilon_{ij}^{(\mathrm{Y})}$ has two identical eigenvalues ($\varepsilon_1^{(\mathrm{Y})} \neq \varepsilon_2^{(\mathrm{Y})} = \varepsilon_3^{(\mathrm{Y})}$) which gives the following analytical expression for $\mathbf{C}^{(\mathrm{F})}$

$$\mathbf{C}^{(\mathrm{F})} = s_1 \frac{\mathrm{d}\left(\boldsymbol{\varepsilon}^{(\mathrm{Y})}\right)^2}{\mathrm{d}\boldsymbol{\varepsilon}^{(\mathrm{Y})}} - s_2 \mathbf{I}_\mathrm{S} - s_3 \boldsymbol{\varepsilon}^{(\mathrm{Y})} \otimes \boldsymbol{\varepsilon}^{(\mathrm{Y})} + s_4 \boldsymbol{\varepsilon}^{(\mathrm{Y})} \otimes \mathbf{I} + s_5 \mathbf{I} \otimes \boldsymbol{\varepsilon}^{(\mathrm{Y})} - s_6 \mathbf{I} \otimes \mathbf{I} \quad \text{(S1.36)}$$

where

$$\mathrm{I}_{ij} = \delta_{ij} \quad \text{(S1.37)}$$

is the second-order identity tensor, and

$$s_1 = \frac{\sigma_a^{(\mathrm{F})} - \sigma_c^{(\mathrm{F})}}{\left(\varepsilon_a^{(\mathrm{Y})} - \varepsilon_c^{(\mathrm{Y})}\right)^2} + \frac{1}{\varepsilon_a^{(\mathrm{Y})} - \varepsilon_c^{(\mathrm{Y})}}\left(\frac{\partial \sigma_c^{(\mathrm{F})}}{\partial \varepsilon_b^{(\mathrm{Y})}} - \frac{\partial \sigma_c^{(\mathrm{F})}}{\partial \varepsilon_c^{(\mathrm{Y})}}\right) \quad \text{(S1.38a)}$$

$$s_2 = 2\varepsilon_c^{(\mathrm{Y})} \frac{\sigma_a^{(\mathrm{F})} - \sigma_c^{(\mathrm{F})}}{\left(\varepsilon_a^{(\mathrm{Y})} - \varepsilon_c^{(\mathrm{Y})}\right)^2} + \frac{\varepsilon_a^{(\mathrm{Y})} + \varepsilon_c^{(\mathrm{Y})}}{\varepsilon_a^{(\mathrm{Y})} - \varepsilon_c^{(\mathrm{Y})}}\left(\frac{\partial \sigma_c^{(\mathrm{F})}}{\partial \varepsilon_b^{(\mathrm{Y})}} - \frac{\partial \sigma_c^{(\mathrm{F})}}{\partial \varepsilon_c^{(\mathrm{Y})}}\right) \quad \text{(S1.38b)}$$

$$s_3 = 2 \frac{\sigma_a^{(\mathrm{F})} - \sigma_c^{(\mathrm{F})}}{\left(\varepsilon_a^{(\mathrm{Y})} - \varepsilon_c^{(\mathrm{Y})}\right)^3} + \frac{1}{\left(\varepsilon_a^{(\mathrm{Y})} - \varepsilon_c^{(\mathrm{Y})}\right)^2}\left(\frac{\partial \sigma_a^{(\mathrm{F})}}{\partial \varepsilon_c^{(\mathrm{Y})}} + \frac{\partial \sigma_c^{(\mathrm{F})}}{\partial \varepsilon_a^{(\mathrm{Y})}} - \frac{\partial \sigma_a^{(\mathrm{F})}}{\partial \varepsilon_a^{(\mathrm{Y})}} - \frac{\partial \sigma_c^{(\mathrm{F})}}{\partial \varepsilon_c^{(\mathrm{Y})}}\right) \quad \text{(S1.38c)}$$

$$s_4 = 2\varepsilon_c^{(\mathrm{Y})} \frac{\sigma_a^{(\mathrm{F})} - \sigma_c^{(\mathrm{F})}}{\left(\varepsilon_a^{(\mathrm{Y})} - \varepsilon_c^{(\mathrm{Y})}\right)^3} + \frac{1}{\varepsilon_a^{(\mathrm{Y})} - \varepsilon_c^{(\mathrm{Y})}}\left(\frac{\partial \sigma_a^{(\mathrm{F})}}{\partial \varepsilon_c^{(\mathrm{Y})}} - \frac{\partial \sigma_c^{(\mathrm{F})}}{\partial \varepsilon_b^{(\mathrm{Y})}}\right) + \frac{\varepsilon_c^{(\mathrm{Y})}}{\left(\varepsilon_a^{(\mathrm{Y})} - \varepsilon_c^{(\mathrm{Y})}\right)^2}\left(\frac{\partial \sigma_a^{(\mathrm{F})}}{\partial \varepsilon_c^{(\mathrm{Y})}} + \frac{\partial \sigma_c^{(\mathrm{F})}}{\partial \varepsilon_a^{(\mathrm{Y})}} - \frac{\partial \sigma_a^{(\mathrm{F})}}{\partial \varepsilon_a^{(\mathrm{Y})}} - \frac{\partial \sigma_c^{(\mathrm{F})}}{\partial \varepsilon_c^{(\mathrm{Y})}}\right) \quad \text{(S1.38d)}$$

$$s_5 = 2\varepsilon_c^{(\mathrm{Y})} \frac{\sigma_a^{(\mathrm{F})} - \sigma_c^{(\mathrm{F})}}{\left(\varepsilon_a^{(\mathrm{Y})} - \varepsilon_c^{(\mathrm{Y})}\right)^3} + \frac{1}{\varepsilon_a^{(\mathrm{Y})} - \varepsilon_c^{(\mathrm{Y})}}\left(\frac{\partial \sigma_c^{(\mathrm{F})}}{\partial \varepsilon_a^{(\mathrm{Y})}} - \frac{\partial \sigma_c^{(\mathrm{F})}}{\partial \varepsilon_b^{(\mathrm{Y})}}\right) + \frac{\varepsilon_c^{(\mathrm{Y})}}{\left(\varepsilon_a^{(\mathrm{Y})} - \varepsilon_c^{(\mathrm{Y})}\right)^2}\left(\frac{\partial \sigma_a^{(\mathrm{F})}}{\partial \varepsilon_c^{(\mathrm{Y})}} + \frac{\partial \sigma_c^{(\mathrm{F})}}{\partial \varepsilon_a^{(\mathrm{Y})}} - \frac{\partial \sigma_a^{(\mathrm{F})}}{\partial \varepsilon_a^{(\mathrm{Y})}} - \frac{\partial \sigma_c^{(\mathrm{F})}}{\partial \varepsilon_c^{(\mathrm{Y})}}\right) \quad \text{(S1.38e)}$$

$$s_6 = 2\varepsilon_c^{(\mathrm{Y})} \frac{\sigma_a^{(\mathrm{F})} - \sigma_c^{(\mathrm{F})}}{\left(\varepsilon_a^{(\mathrm{Y})} - \varepsilon_c^{(\mathrm{Y})}\right)^3} + \frac{\varepsilon_a^{(\mathrm{Y})}\varepsilon_c^{(\mathrm{Y})}}{\left(\varepsilon_a^{(\mathrm{Y})} - \varepsilon_c^{(\mathrm{Y})}\right)^2}\left(\frac{\partial \sigma_a^{(\mathrm{F})}}{\partial \varepsilon_c^{(\mathrm{Y})}} + \frac{\partial \sigma_c^{(\mathrm{F})}}{\partial \varepsilon_a^{(\mathrm{Y})}}\right) - \frac{\left(\varepsilon_c^{(\mathrm{Y})}\right)^2}{\left(\varepsilon_a^{(\mathrm{Y})} - \varepsilon_c^{(\mathrm{Y})}\right)^2}\left(\frac{\partial \sigma_a^{(\mathrm{F})}}{\partial \varepsilon_a^{(\mathrm{Y})}} + \frac{\partial \sigma_c^{(\mathrm{F})}}{\partial \varepsilon_c^{(\mathrm{Y})}}\right) - \frac{\varepsilon_a^{(\mathrm{Y})} + \varepsilon_c^{(\mathrm{Y})}}{\varepsilon_a^{(\mathrm{Y})} - \varepsilon_c^{(\mathrm{Y})}} \frac{\partial \sigma_c^{(\mathrm{F})}}{\partial \varepsilon_b^{(\mathrm{Y})}} \quad \text{(S1.38f)}$$

are constants with $(a, b, c)$ being cyclic permutations of (1,2,3). Finally, In the third case, the three eigenvalues of the strain tensor $\varepsilon_{ij}^{(\mathrm{Y})}$ are all identical ($\varepsilon_1^{(\mathrm{Y})} = \varepsilon_2^{(\mathrm{Y})} = \varepsilon_3^{(\mathrm{Y})}$) which gives the following analytical expression for $\mathbf{C}^{(\mathrm{F})}$

$$\mathbf{C}^{(\mathrm{F})} = \left(\frac{\partial \sigma_1^{(\mathrm{F})}}{\partial \varepsilon_1^{(\mathrm{Y})}} - \frac{\partial \sigma_1^{(\mathrm{F})}}{\partial \varepsilon_2^{(\mathrm{Y})}}\right)\mathbf{I}_\mathrm{S} + \frac{\partial \sigma_1^{(\mathrm{F})}}{\partial \varepsilon_2^{(\mathrm{Y})}}\mathbf{I} \otimes \mathbf{I} \tag{S1.39}$$

Note that all three cases require $\partial \sigma_i^{(\mathrm{F})} / \partial \varepsilon_j^{(\mathrm{Y})}$ which can be determined by taking the first derivative of $\sigma_i^{(\mathrm{F})}$ in (S1.30)

$$\frac{\partial \sigma_i^{(\mathrm{F})}}{\partial \varepsilon_i^{(\mathrm{Y})}} = \frac{\partial}{\partial \varepsilon_i^{(\mathrm{Y})}}\left(\frac{\partial f}{\partial \varepsilon_i^{(\mathrm{Y})}}\right) = \begin{cases} 0 & \varepsilon_i^{(\mathrm{Y})} < \epsilon_1 \\ \ell \dfrac{\left(\dfrac{\varepsilon_i^{(\mathrm{Y})} - \epsilon_1}{\epsilon_2 - \epsilon_1}\right)^t \left(\varepsilon_i^{(\mathrm{Y})} - \epsilon_1\right)}{t+1} & \epsilon_1 \le \varepsilon_i^{(\mathrm{Y})} < \epsilon_2 \\ \ell \left[\dfrac{\left(1 + \varepsilon_i^{(\mathrm{Y})} - \epsilon_2\right)^{s+1} - 1}{s+1} + \dfrac{\epsilon_2 - \epsilon_1}{t+1}\right] & \varepsilon_i^{(\mathrm{Y})} \ge \epsilon_2 \end{cases} \tag{S1.40}$$

With $\mathbf{C}^{(\mathrm{F})}$ at hand, the stiffness of actin filament network $\mathbf{C}^{(\mathrm{A})}$ can be obtained from equation (S1.21).

#### 1.2. Total cell stiffness

Next, we determine the total stiffness of the cell. To this end, we first degrade the fourth-order tensors $\mathbf{C}^{(\mathrm{MT})}$ (S1.16), $\mathbf{C}^{(\rho)}$ (S1.19), and $\mathbf{C}^{(\mathrm{A})}$ (S1.21) to the second-order tensors $\mathbf{C}^{(\mathrm{MT})}$, $\mathbf{C}^{(\rho)}$, and $\mathbf{C}^{(\mathrm{A})}$. Equation (1.20) illustrates how a fourth-order tensor (*e.g.*, $\mathrm{C}_{ijkl}^{(\rho)}$) can be degraded to a 6×6 square matrix (*e.g.*, $\mathrm{C}_{ij}^{(\rho)}$). As the actin filament network in our model is connected to the myosin motors and the microtubules in series (Figure S18), the total stiffness of the cell, $\mathbf{C}$, is obtained as follows

$$\mathbf{C} = \left(\left(\mathbf{C}^{(\mathrm{X})}\right)^{-1} + \left(\mathbf{C}^{(\mathrm{Y})}\right)^{-1}\right)^{-1} \tag{S1.41}$$

where

$$\mathbf{C}^{(\mathrm{X})} = \mathbf{C}^{(\rho)} + \mathbf{C}^{(\mathrm{MT})} \tag{S1.42}$$

and

$$\mathbf{C}^{(\mathrm{Y})} = \mathbf{C}^{(\mathrm{A})} = \mathbf{C}^{(\mathrm{I})} + \mathbf{C}^{(\mathrm{F})} \tag{S1.43}$$

#### 1.3. Solving the set of nonlinear equations

In the previous sections, we presented the constitutive equations of the cell model where the stress field $\sigma_{ij}$ can be determined from (S1.17) or (S1.25) and the stiffness field $\mathrm{C}_{ij}$ can be obtained from equation (S1.41). However, $\sigma_{ij}$ and $\mathrm{C}_{ij}$ are functions of the strain tensors $\varepsilon_{ij}^{(\mathrm{X})}$ and $\varepsilon_{ij}^{(\mathrm{Y})}$ (and not $\varepsilon_{ij}$) which are both unknown. To determine the unknown tensors $\varepsilon_{ij}^{(\mathrm{X})}$ and $\varepsilon_{ij}^{(\mathrm{Y})}$, we first define the following 12×1 vector

$$\begin{aligned}\boldsymbol{u} = &\left\{\varepsilon_{11}^{(\mathrm{X})} \quad \varepsilon_{22}^{(\mathrm{X})} \quad \varepsilon_{33}^{(\mathrm{X})} \quad \varepsilon_{12}^{(\mathrm{X})} \quad \varepsilon_{13}^{(\mathrm{X})} \quad \varepsilon_{23}^{(\mathrm{X})} \quad \varepsilon_{11}^{(\mathrm{Y})} \quad \varepsilon_{22}^{(\mathrm{Y})} \quad \varepsilon_{33}^{(\mathrm{Y})} \quad \varepsilon_{12}^{(\mathrm{Y})} \quad \varepsilon_{13}^{(\mathrm{Y})} \quad \varepsilon_{23}^{(\mathrm{Y})}\right\}^\mathrm{T} \\ &= \{u_1 \quad u_2 \quad \cdots \quad u_{12}\}^\mathrm{T}\end{aligned} \tag{S1.44}$$

which contains all 12 unknown variables in the tensors $\varepsilon_{ij}^{(\mathrm{X})}$ and $\varepsilon_{ij}^{(\mathrm{Y})}$. To determine the 12 unknowns, we need 12 equations where 6 of them can be obtained from the following condition

$$\boldsymbol{\sigma} = \boldsymbol{\sigma}^{(\mathrm{X})} = \boldsymbol{\sigma}^{(\mathrm{Y})} \tag{S1.45}$$

where the stress generated by the actomyosin contractility $\boldsymbol{\sigma}^{(X)}$ (equations (S1.17) and (S1.18))

$$\sigma_{ij}^{(X)} = \sigma_{ij} = \left(C_{ijkl}^{(\rho)} + C_{ijkl}^{(MT)}\right)\varepsilon_{kl}^{(X)} + \bar{\rho}_0\delta_{ij} \tag{S1.46}$$

is directly transmitted to the actin filament network $\boldsymbol{\sigma}^{(Y)}$ (equation (S1.41))

$$\sigma_{ij}^{(Y)} = \sigma_{ij} = \sigma_{ij}^{(A)} = \sigma_{ij}^{(I)} + \sigma_{ij}^{(F)} \tag{S1.47}$$

The other 6 equations are given as follows

$$\boldsymbol{\varepsilon} = \boldsymbol{\varepsilon}^{(X)} + \boldsymbol{\varepsilon}^{(Y)} \tag{S1.48}$$

where $\boldsymbol{\varepsilon}^{(X)}$ is the strain of the cytoskeletal components that are in compression (*e.g.*, microtubule network), $\boldsymbol{\varepsilon}^{(Y)}$ is the strain of the cytoskeletal components that are in tension (*e.g.*, actin filaments), and $\boldsymbol{\varepsilon}$ is the total strain of the cell. As all stress and strain tensors $\sigma_{ij}^{(X)}$, $\sigma_{ij}^{(Y)}$, $\varepsilon_{ij}^{(X)}$, and $\varepsilon_{ij}^{(Y)}$ are symmetric, the conditions in (S1.45) and (S1.48) can be defined by the following 12 equations in the 12×1 vector $\boldsymbol{f}$

$$\boldsymbol{f} = \{f_1 \quad f_2 \quad \cdots \quad f_{12}\}^{T} \tag{S1.49a}$$

where

$$f_1 = \sigma_{11}^{(X)} - \sigma_{11}^{(Y)} \tag{S1.49b}$$
$$f_2 = \sigma_{22}^{(X)} - \sigma_{22}^{(Y)} \tag{S1.49c}$$
$$f_3 = \sigma_{33}^{(X)} - \sigma_{33}^{(Y)} \tag{S1.49d}$$
$$f_4 = \sigma_{12}^{(X)} - \sigma_{12}^{(Y)} \tag{S1.49e}$$
$$f_5 = \sigma_{13}^{(X)} - \sigma_{13}^{(Y)} \tag{S1.49f}$$
$$f_6 = \sigma_{23}^{(X)} - \sigma_{23}^{(Y)} \tag{S1.49g}$$
$$f_7 = \varepsilon_{11} - \varepsilon_{11}^{(X)} - \varepsilon_{11}^{(Y)} \tag{S1.49h}$$
$$f_8 = \varepsilon_{22} - \varepsilon_{22}^{(X)} - \varepsilon_{22}^{(Y)} \tag{S1.49i}$$
$$f_9 = \varepsilon_{33} - \varepsilon_{33}^{(X)} - \varepsilon_{33}^{(Y)} \tag{S1.49j}$$
$$f_{10} = \varepsilon_{12} - \varepsilon_{12}^{(X)} - \varepsilon_{12}^{(Y)} \tag{S1.49k}$$
$$f_{11} = \varepsilon_{13} - \varepsilon_{13}^{(X)} - \varepsilon_{13}^{(Y)} \tag{S1.49l}$$
$$f_{12} = \varepsilon_{23} - \varepsilon_{23}^{(X)} - \varepsilon_{23}^{(Y)} \tag{S1.49m}$$

We then determine the 12×12 Jacobian matrix $\mathbf{J}$

$$\mathbf{J} = \begin{bmatrix} \partial f_1/\partial u_1 & \partial f_1/\partial u_2 & \cdots & \partial f_1/\partial u_{12} \\ \partial f_2/\partial u_1 & \partial f_2/\partial u_2 & \cdots & \partial f_2/\partial u_{12} \\ \vdots & \vdots & & \vdots \\ \partial f_{12}/\partial u_1 & \partial f_{12}/\partial u_2 & \cdots & \partial f_{12}/\partial u_{12} \end{bmatrix} = \begin{bmatrix} \mathbf{C}^{(X)} & -\mathbf{C}^{(Y)} \\ -\mathbf{I} & -\mathbf{I} \end{bmatrix} \tag{S1.50}$$

using equations (S1.44) and (S1.49) for $u_i$ and $f_i$, respectively. Finally, we use the Newton-Raphson method

$$\boldsymbol{u}_{i+1} = \boldsymbol{u}_i - \mathbf{J}^{-1}\boldsymbol{f}(\boldsymbol{u}_i) \tag{S1.51}$$

to determine the unknown vector $\boldsymbol{u}$ where the vectors $\boldsymbol{u}_i$ and $\boldsymbol{u}_{i+1}$ are respectively the solutions for $i$ and $i+1$ iterations. To find the solutions of the nonlinear equations, we use the following initial guess $\boldsymbol{u}_0$ (or any other initial guess)

$$\boldsymbol{u}_0 = \{0 \quad 0 \quad \ldots \quad 0\}_{1\times 12}^{T} \tag{S1.52}$$

and the convergence criterion

$$|\boldsymbol{f}| = \sqrt{(f_1)^2 + (f_2)^2 + \cdots + (f_{12})^2} < \epsilon_{\mathrm{Tol}} \tag{S1.53}$$

where $|\boldsymbol{f}|$ is the magnitude of the vector $\boldsymbol{f}$, and $\epsilon_{\mathrm{Tol}}$ is the convergence threshold. Using $\epsilon_{\mathrm{Tol}} = 10^{-8}$ in our simulations, we stop the iterations in (S1.51) when $|\boldsymbol{f}|$ is less than $\epsilon_{\mathrm{Tol}}$. With the strain tensors $\varepsilon_{ij}^{(X)}$ and $\varepsilon_{ij}^{(Y)}$ determined from (S1.51), we can calculate the stress tensor $\sigma_{ij}$ (from (S1.17) or (S1.25)) and the stiffness tensor $C_{ij}$ (from (S1.41)).

#### 1.4. Adding the vimentin network to the model

After describing the model details, we here show how the intermediate filament network can be included in the model. As described above, both stress tensor $\sigma_{ij}$ and stiffness tensor $\mathrm{C}_{ij}$ should be defined to have a complete set of constitutive equations. The stress tensor $\sigma_{ij}$ can be obtained from either (S1.17) or (S1.25), and the stiffness tensor $\mathrm{C}_{ij}$ is calculated by (S1.41). We here describe how $\sigma_{ij}$ and $\mathrm{C}_{ij}$ should be modified to include vimentin filaments in the model.

In the main text, it was described that the model has two vimentin elements. The first element is parallel with the actin element and experiences tensile stresses, while the second element is parallel with the microtubule element and undergoes compression (Figure S18). To include the tensile vimentin element in $\sigma_{ij}$, equation (S1.25) should be rewritten as follows

$$\sigma_{ij} = \sigma_{ij}^{(\mathrm{I})} + \sigma_{ij}^{(\mathrm{F})} + \sigma_{ij}^{(\mathrm{VT})} \quad (\mathrm{S1.54})$$

where $\boldsymbol{\sigma}^{(\mathrm{VT})}$

$$\sigma_{ij}^{(\mathrm{VT})} = \mathrm{C}_{ijkl}^{(\mathrm{VT})}\,\varepsilon_{kl}^{(\mathrm{Y})} \quad (\mathrm{S1.55})$$

is linearly related to the strain tensor $\boldsymbol{\varepsilon}^{(\mathrm{Y})}$ using the stiffness tensor $\mathbf{C}^{(\mathrm{VT})}$

$$\mathrm{C}_{ijkl}^{(\mathrm{VT})} = K^{(\mathrm{VT})}\delta_{ij}\delta_{kl} + \mu^{(\mathrm{VT})}\left(\delta_{ik}\delta_{jk} + \delta_{il}\delta_{jk} - \frac{2}{3}\delta_{ij}\delta_{kl}\right) \quad (\mathrm{S1.56})$$

where

$$K^{(\mathrm{VT})} = \frac{E^{(\mathrm{VT})}}{3(1-2v^{(\mathrm{VT})})} \quad (\mathrm{S1.57})$$

is the bulk modulus of the tensile vimentin network, and

$$\mu^{(\mathrm{VT})} = \frac{E^{(\mathrm{VT})}}{2(1+v^{(\mathrm{VT})})} \quad (\mathrm{S1.58})$$

is the shear modulus of the tensile vimentin network, $E^{(\mathrm{VT})}$ is the elastic modulus of the tensile vimentin network, and $v^{(\mathrm{VT})}$ is the Poisson's ratio of the tensile vimentin network. Note that in (S1.55), tensile vimentin filaments are assumed to behave linearly. However, (S1.55) can be replaced by a nonlinear equation to have a more general form and to account for the tension-stiffening of vimentin filaments. Similarly, to include the compressive vimentin element in $\sigma_{ij}$, equation (S1.17) should be rewritten as follows

$$\rho_{ij} = -\mathrm{C}_{ijkl}^{(\mathrm{MT})}\,\varepsilon_{kl}^{(\mathrm{X})} - \mathrm{C}_{ijkl}^{(\mathrm{VC})}\,\varepsilon_{kl}^{(\mathrm{X})} + \sigma_{ij} \quad (\mathrm{S1.59})$$

where

$$\mathrm{C}_{ijkl}^{(\mathrm{VC})} = K^{(\mathrm{VC})}\delta_{ij}\delta_{kl} + \mu^{(\mathrm{VC})}\left(\delta_{ik}\delta_{jk} + \delta_{il}\delta_{jk} - \frac{2}{3}\delta_{ij}\delta_{kl}\right) \quad (\mathrm{S1.60})$$

is the stiffness tensor of the compressive vimentin element and

$$K^{(\mathrm{VC})} = \frac{E^{(\mathrm{VC})}}{3(1-2v^{(\mathrm{VC})})} \quad (\mathrm{S1.61})$$

is the bulk modulus of the compressive vimentin network, and

$$\mu^{(\mathrm{VC})} = \frac{E^{(\mathrm{VC})}}{2(1+v^{(\mathrm{VC})})} \quad (\mathrm{S1.62})$$

is the shear modulus of the compressive vimentin network, $E^{(\mathrm{VC})}$ is the elastic modulus of the compressive vimentin network, and $v^{(\mathrm{VC})}$ is the Poisson's ratio of the compressive vimentin network.

Finally, to include the compressive and tensile vimentin elements in $\mathrm{C}_{ij}$, the total stiffness of the cell should be rewritten as follows

$$\mathbf{C} = \left(\left(\mathbf{C}^{(\mathrm{X})}\right)^{-1} + \left(\mathbf{C}^{(\mathrm{Y})}\right)^{-1}\right)^{-1} \quad (\mathrm{S1.63})$$

where

$$\mathbf{C}^{(\mathrm{X})} = \mathbf{C}^{(\rho)} + \mathbf{C}^{(\mathrm{MT})} + \mathbf{C}^{(\mathrm{VC})} \tag{S1.64}$$

and

$$\mathbf{C}^{(\mathrm{Y})} = \mathbf{C}^{(\mathrm{A})} = \mathbf{C}^{(\mathrm{I})} + \mathbf{C}^{(\mathrm{F})} + \mathbf{C}^{(\mathrm{VT})} \tag{S1.65}$$

Note that the stress tensors $\sigma_{ij}^{(\mathrm{c})}$ and $\sigma_{ij}^{(\mathrm{t})}$ described in the main text (and their eigenvalues shown in Figure 5) represent the compressive and tensile stresses that the compressive (parallel) and tensile (series) elements of the model experience due to the cell contractility $\rho_{ij}$, respectively,

$$\sigma_{ij}^{(\mathrm{c})} = \mathrm{C}_{ijkl}^{(\mathrm{MT})}\varepsilon_{kl}^{(\mathrm{X})} + \mathrm{C}_{ijkl}^{(\mathrm{VC})}\varepsilon_{kl}^{(\mathrm{X})} \tag{S1.66}$$

$$\sigma_{ij}^{(\mathrm{t})} = \sigma_{ij} = \sigma_{ij}^{(\mathrm{I})} + \sigma_{ij}^{(\mathrm{F})} + \sigma_{ij}^{(\mathrm{VT})} \tag{S1.67}$$

#### 2. One-dimensional cell model

We here describe a one-dimensional framework of the cell model to present the key features of the model without the complexity of the three-dimensional framework. As described in the main text, the model is composed of the following elements: (i) the myosin molecular motors, (ii) the microtubules, (iii) the actin filaments, and (iv) the vimentin filaments. We first derive the constitutive equations of the model without the presence of the vimentin element and we will later describe how the vimentin filament network can be added to the model.

Phosphorylated myosin molecular motors generate internal stresses which are denoted by $\rho$ (cell contractility) in our one-dimensional model and is defined as follows (*7*)

$$\rho = \frac{E^{(\mathrm{MT})}\,\alpha - 1}{\beta - \alpha}\varepsilon^{(\mathrm{X})} + \frac{\beta}{\beta - \alpha}\rho_0 \tag{S2.1}$$

where $\alpha$ is the chemo-mechanical feedback parameter, $\beta$ is the chemical stiffness parameter, $\varepsilon^{(\mathrm{X})}$ is the strain of the compressive elements shown in Figure S18, $\rho_0$ is the initial contractility of the cell, and $E^{(\mathrm{MT})}$ is the stiffness of the microtubule network. A large value of $\alpha$ strengthens the stress-dependent feedback mechanism of the cell and the cell promotes myosin motor binding in response to the cytoskeletal tension. Subsequently, this increase in the overall density of phosphorylated myosin molecular motors increases the cell contractility $\rho$, the cell generated active stress $\sigma$, the cytoskeletal stiffness $E$, and the cell strain $|\varepsilon|$. Unlike $\alpha$, a large value of $\beta$ weakens the stress-dependent feedback mechanism of the cell as it makes myosin motor recruitment more difficult. Therefore, an increase in $\beta$ causes cell contractility, active stress, cytoskeletal stiffness, and cell strain to decrease.

Microtubules are compressively loaded by the cell-generated internal stress while the rest of this internal stress is transmitted to the extracellular matrix through the actin filament network. Denoting the stress in the microtubule network as $E^{(\mathrm{MT})}\varepsilon^{(\mathrm{X})}$ and using $\sigma$ to denote the stress transmitted to the extracellular matrix through the actin filament network, we have

$$\rho = -E^{(\mathrm{MT})}\varepsilon^{(\mathrm{X})} + \sigma \tag{S2.2}$$

Substituting equation (S2.1) into (S2.2), we can derive the tensile stress $\sigma$ generated by the cell

$$\sigma = \frac{E^{(\mathrm{MT})}\,\beta - 1}{\beta - \alpha}\varepsilon^{(\mathrm{X})} + \frac{\beta}{\beta - \alpha}\rho_0 \tag{S2.3}$$

which is transmitted to the extracellular matrix through the actin filament network

$$\sigma = E^{(\mathrm{A})}\varepsilon^{(\mathrm{Y})} \tag{S2.4}$$

where $E^{(\mathrm{A})}$ is the stiffness of the actin network, and $\varepsilon^{(\mathrm{Y})}$ is the strain of the tensile elements shown in Figure S18. Consistent with experimental observations (*5*), the stiffness of the actin network $E^{(\mathrm{A})}$ increases in response to the cytoskeletal tension $\sigma$. To capture the stiffening of the actin network, $E^{(\mathrm{A})}$ increases with tension (but not in compression) for tensile stresses beyond a critical tensile stress

$$\begin{cases} E^{(\mathrm{A})} = E^{(\mathrm{I})} & \varepsilon_i^{(\mathrm{Y})} < \epsilon_{\mathrm{A}} \\ E^{(\mathrm{A})} = E^{(\mathrm{I})} + E^{(\mathrm{I})} \ell (\varepsilon^{(\mathrm{Y})} - \epsilon_{\mathrm{A}})^m & \varepsilon_i^{(\mathrm{Y})} \geq \epsilon_{\mathrm{A}} \end{cases} \quad (S2.5)$$

where $E^{(\mathrm{I})}$ is the initial elastic modulus of the actin network (*e.g.*, the elastic modulus of the actin network when the stiffness of the matrix is negligible $\rightarrow E^{(\mathrm{A})} = E^{(\mathrm{I})}$), and $\ell$ and $m$ are the stiffening parameters. Either of these stiffening parameters can regulate the stiffening of the actin network. Therefore, to reduce the number of fitting parameters, we set $\ell$ = 200 and we only consider $m$ as a fitting parameter in the model.

Assuming that the matrix exhibits a linear elastic behavior

$$\sigma = E^{(\mathrm{m})} \varepsilon^{(\mathrm{m})} \quad (S2.6)$$

the cell-generated matrix strain $\varepsilon^{(\mathrm{m})}$ can be determined as follows

$$2\varepsilon^{(\mathrm{m})} + \varepsilon = 2\varepsilon^{(\mathrm{m})} + \varepsilon^{(\mathrm{X})} + \varepsilon^{(\mathrm{Y})} = 0 \quad (S2.7)$$

Solving equations (S2.3), (S2.4), (S2.6), and (S2.7) together, we can determine $\sigma$, $\varepsilon^{(\mathrm{X})}$, $\varepsilon^{(\mathrm{Y})}$, and $\varepsilon^{(\mathrm{m})}$ (4 equations, 4 unknowns). Then, the cell contractility $\rho$ can be obtained from (S2.1) or (S2.2). Note that $\beta$ and $\alpha$ cannot be equal because for $\beta = \alpha$ the cell contractility $\rho$ and the stress $\sigma$ become infinity in equations (S2.1) and (S2.3), respectively, which are not physically possible. Also, $E^{(\mathrm{MT})}$ should be always positive as the elastic modulus of the microtubule network cannot be negative. Furthermore, the cell length should decrease with cell contraction which yields $\varepsilon^{(\mathrm{X})} \leq 0$. Finally, to ensure that $\rho$ is always positive (i.e., actomyosin contractility pulls on, and not pushes, the extracellular matrix), the two coefficients of $(E^{(\mathrm{MT})} \alpha - 1)/(\beta - \alpha)$ and $\beta/(\beta - \alpha)$ in (S2.1) should be positive. These constraints give the following stability criterion

$$\beta > \alpha > \frac{1}{E^{(\mathrm{MT})}} \quad (S2.8)$$

After describing the model stability criterion and details, we here show how the intermediate filament network can be included in the model. As described in the main text and shown in Figure S18, the model has two vimentin elements. The first element is parallel with the actin element and experiences tensile stresses, while the second element is parallel with the microtubule element and undergoes compression. To include the compressive vimentin element, $E^{(\mathrm{MT})}$ in the above equations should be replaced by $E^{(\mathrm{MT})} + E^{(\mathrm{VC})}$ where $E^{(\mathrm{VC})}$ is the stiffness of the compressive vimentin network. Similarly, to include the tensile vimentin element, $E^{(\mathrm{A})}$ should be replaced by $E^{(\mathrm{A})} + E^{(\mathrm{VT})}$ where $E^{(\mathrm{VT})}$ is the stiffness of the tensile vimentin network. Note that $E^{(\mathrm{VC})}$ and $E^{(\mathrm{VT})}$ can be constant parameters or they can nonlinearly increase with compression and tension, respectively, to include the strain-stiffening of vimentin filaments in the model. Supplementary Table 1 shows the parameters used in the model. The same set of parameters was used for both 1D and 3D simulations. The model parameters in Supplementary Table 1 were determined by fitting the model to Traction Force Microscopy (TFM) experiments for micropatterned fibroblasts cultured on fibronectin-coated deformable substrates with different stiffness (2.8-30 kPa) and surface areas (700-2400 µm$^2$) (*8*).

#### Supplementary figures

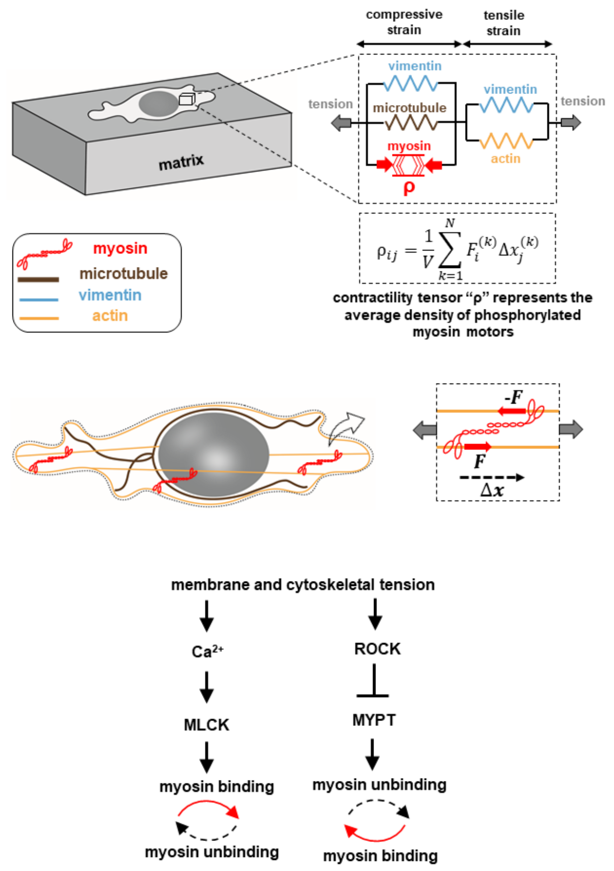


Figure S1. The first component of the cytoskeletal model is myosin which generates internal forces. Phosphorylated myosin motors are represented by active force-generating dipoles $F$. We treat the average density of the force dipoles as a symmetric tensor, $\rho_{ij}$, whose components represent cell contractility in different directions. Various experimental studies show that cell contractility increases with tension through tension-activated signaling pathways such as the Rho-Rock and the $Ca^{2+}$ pathways (*9*–*12*). We, therefore, assume that the average of contractility in all three directions, $\frac{1}{3}\rho_{kk} = (\rho_{11} + \rho_{22} + \rho_{33})/3$, increases with the average of cytoskeletal tension $\frac{1}{3}\sigma_{kk} = (\sigma_{11} + \sigma_{22} + \sigma_{33})/3$, which in turn generates higher cytoskeletal tension. We will later show how the cell contractility $\rho_{ij}$ changes matrix stiffness as a result of this feedback mechanism.

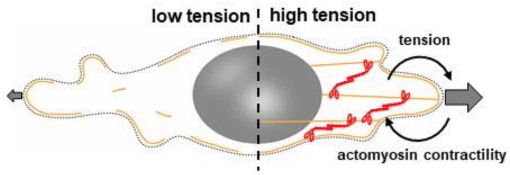


Figure S2. As demonstrated in Figure S1, the Ca$^{+2}$ pathway and the ROCK pathway enable cells to respond to tensile stresses generated at the cell-matrix interface by (i) increasing their contractile forces through increased levels of phosphorylated myosin motors, and (ii) stiffening of the cytoskeleton through recruitment and alignment of actin filaments along the direction of the tensile stresses (see Figure S4). This increase in cell actomyosin contractility, in turn, generates higher tension at the cell-matrix interface leading to a positive feedback loop between actomyosin contractility and cytoskeletal tension.

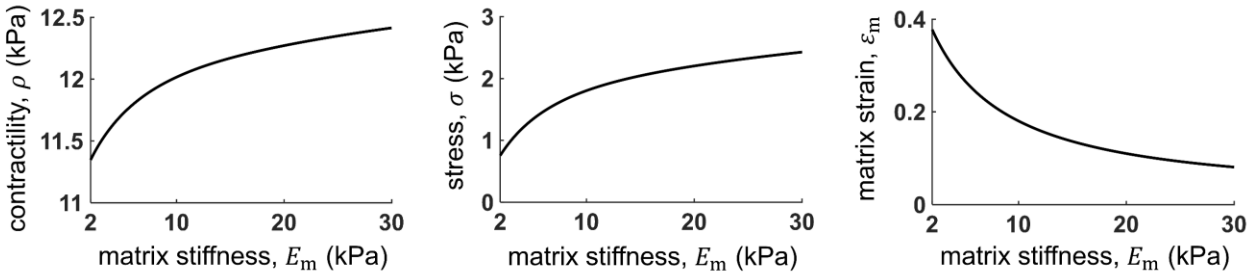


Figure S3. The cell contractility $\rho$, the cell-generated stress $\sigma$, and the cell-generated matrix strain $\varepsilon_{\mathrm{m}}$ as functions of the matrix stiffness $E_{\mathrm{m}}$.

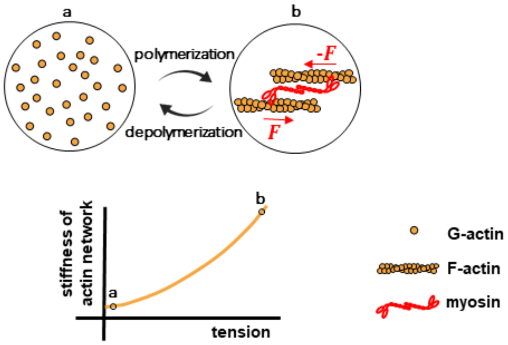


Figure S4. The actin element in the model stiffens with tension, representing the formation of actin filaments and stress fibers in response to tension as observed in experiments (*13*, *14*)

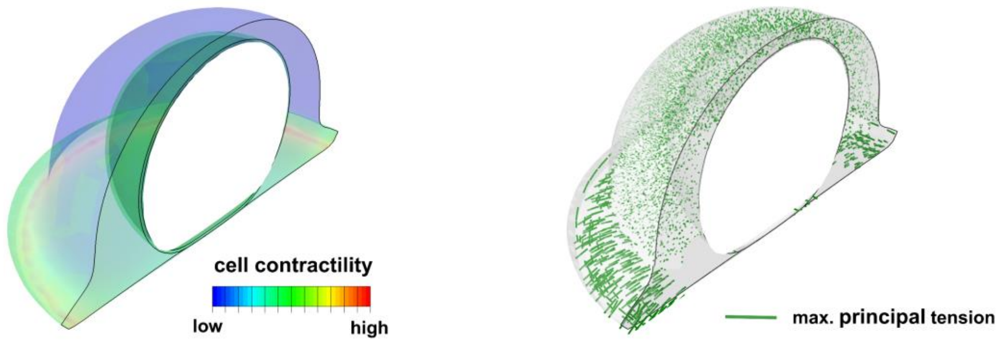


Figure S5. Colocalization of phosphorylated myosin motors and polymerized actin filaments. We simulate cells cultured on a micropatterned substrate with a circular shape (see (*6*) for more details). The cell contractility $\rho_{ij}$ and the actin network stiffness $C_{ijkl}^{(A)}$ in our simulations are initially isotropic (independent of direction) and uniform (independent of spatial location). In other words, cell contractility and actin network stiffness are initially the same everywhere in the cytoplasm with no preferential alignment of phosphorylated myosin motor dipoles and actin filaments. Starting with these initial conditions, our simulations show higher cell contractility in basal regions (compared with apical regions) and close to the cell boundary which is consistent with experimental observations (*6*, *15*, *16*). Concomitant with the increased contractility, our simulations show higher cytoskeletal tension in basal regions close to the cell boundary, which results in stiffening of the actin filaments representing formation of actin filaments as observed experimentally [8].

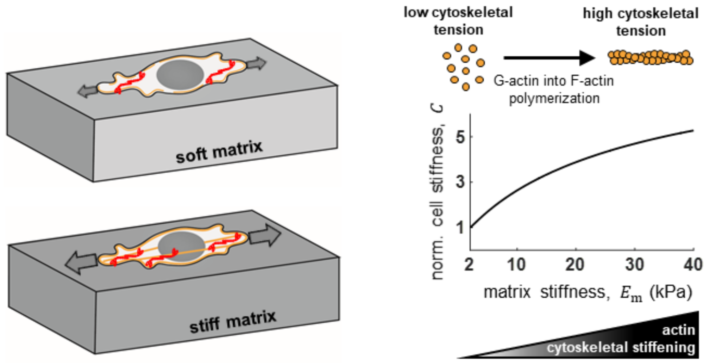


Figure S6. In adherent cells, cellular contraction is resisted by the surrounding matrix leading to generation of mechanical tension at the cell-matrix interface. As matrices with higher stiffness exhibit higher resistance against cellular contraction, cells experience higher tension on these matrices. The increased tension, in turn, activates the $Ca^{+2}$ pathway and the ROCK pathway which enable cells to increase their contractile forces and to stiffen the cytoskeleton through recruitment and alignment of actin filaments along the direction of the tensile stresses.

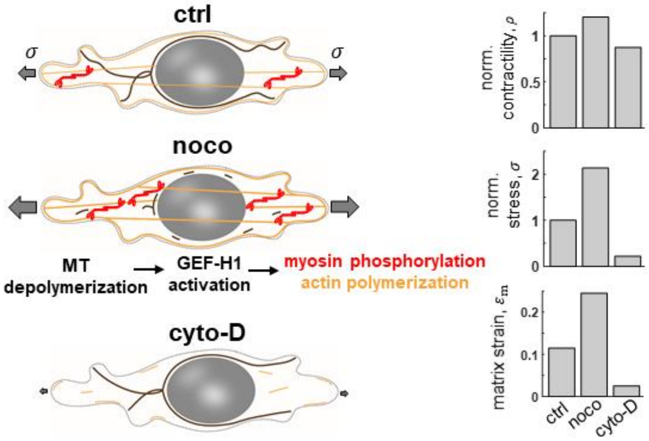


Figure S7. Disruption of the microtubule network increases cell contractility which in turn generates higher tensile stresses, leading to more stretching of the matrix. In contrast, disruption of the actin filament network decreases actomyosin contractility resulting in relaxation of the matrix.

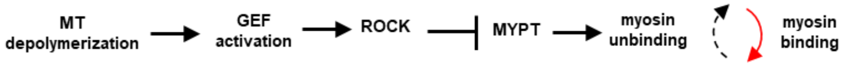


Figure S8. Depolymerization of microtubules activates the ROCK pathway which in turn increases cell contractility and cell force generation (*17*–*21*).

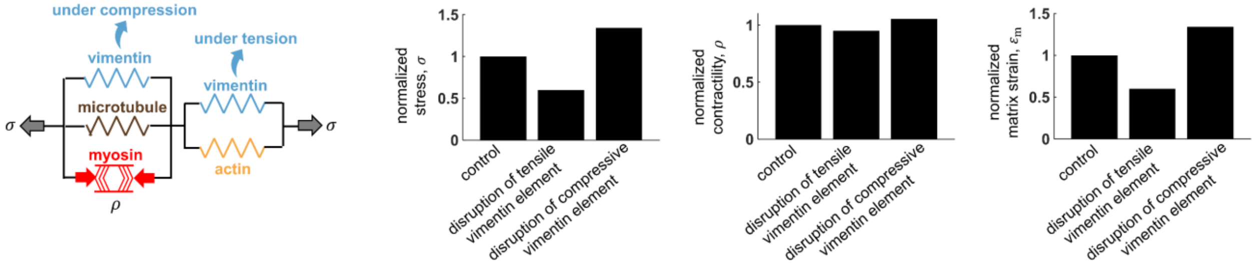


Figure S9. Disruption of the vimentin element in tension (compression) leads to decreases (increases) in the cell contractility $\rho$, the cell-generated stress $\sigma$, and the matrix strain $\varepsilon_{\mathrm{m}}$.

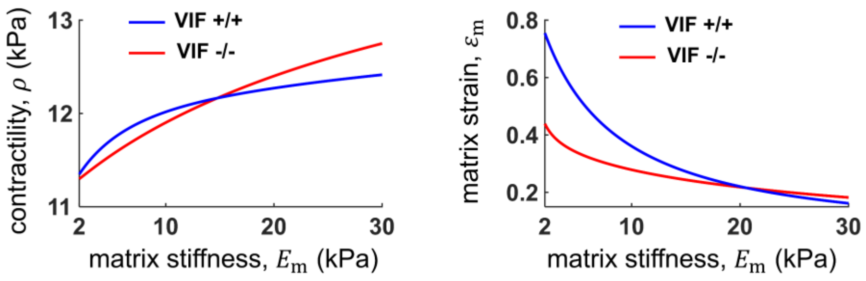


Figure S10. The cell contractility $\rho$ and the cell-generated matrix strain $\varepsilon_m$ as functions of the matrix stiffness $E_m$ with (VIF +/+) and without (VIF -/-) the presence of vimentin intermediate filaments.

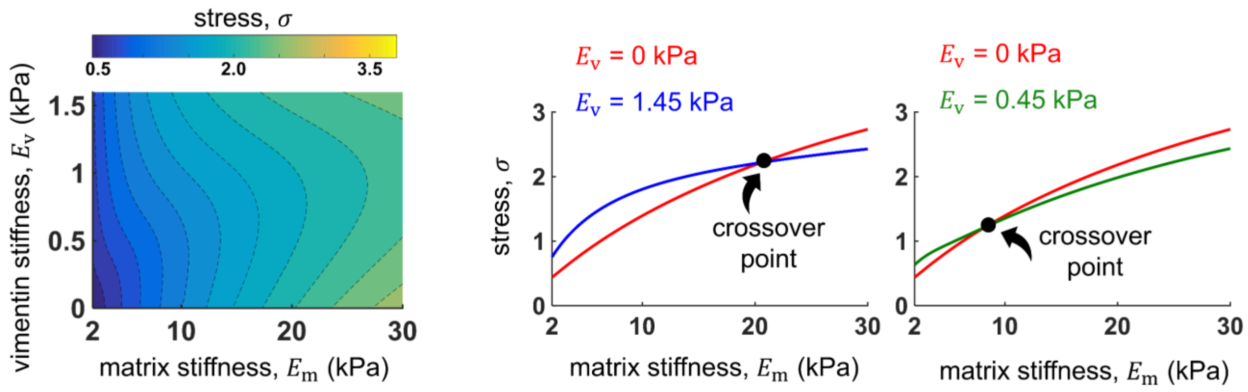


Figure S11. The matrix stiffness at which the crossover occurs depends on the stiffness of the intermediate filaments network. $E_v$ = 0 represents vimentin null cells (VIF -/-)

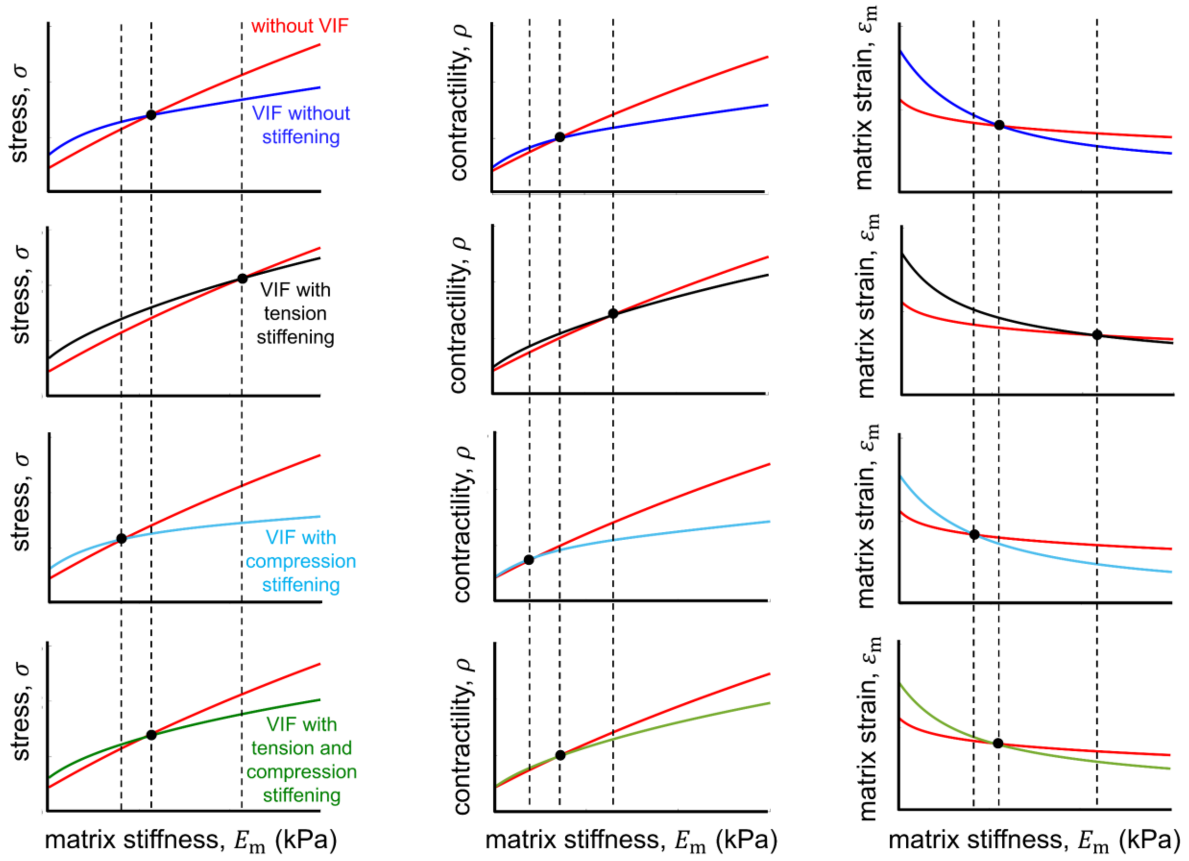


Figure S12. Tension or compression stiffening of intermediate filaments can shift the crossover point to higher or lower matrix stiffness, respectively.

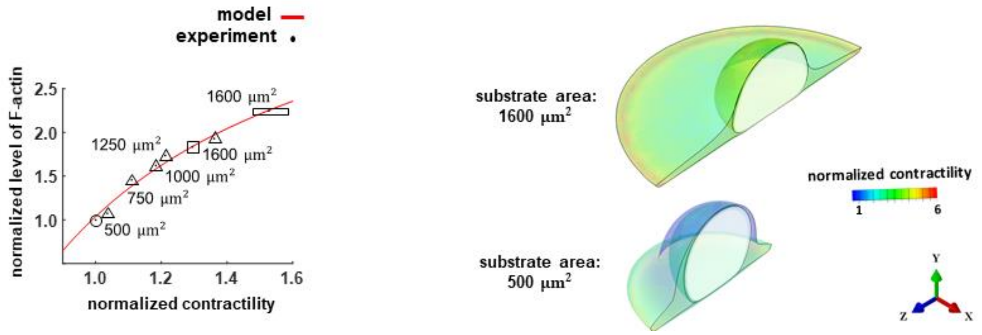


Figure S13. The left panel presents the levels of F-actin and phosphorylated myosin in fibroblasts cultured on rigid micropattern substrates with different substrate areas and shapes. Both F-actin and myosin phosphorylation levels are indicators of cell actomyosin contractility level. Results show that fibroblasts with higher areas are more contractile. For example, fibroblasts cultured on a triangular shape with a substrate area of 1600 μm$^2$ have significantly higher levels of F-actin and myosin than fibroblasts on the same shape with a substrate area of 500 μm$^2$. The right panel shows the cell contractility level spatially within cells cultured on circular substrates with 500 μm$^2$ and 500 μm2 areas (*6*).

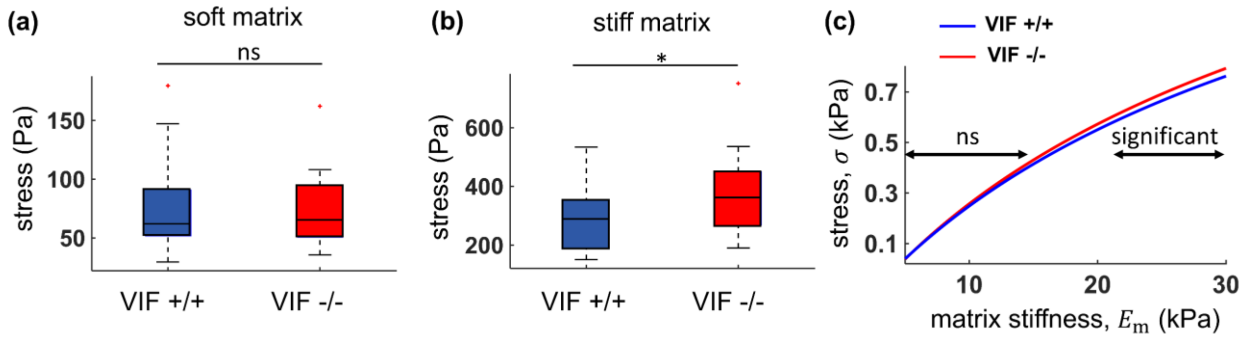


Figure S14. In addition to fibronectin-coated substrates (Figure 3), we also experimentally measure cell-generated traction forces on collagen-coated substrates with low (4.5 kPa) and high (40 kPa) stiffness. Generally, compared with fibronectin-coated substrates, fibroblasts spread less on collagen-coated substrates and generate lower forces, particularly on soft substrates. As a result, the difference between forces generated by VIF -/- and VIF +/+ cells may not be significant on soft substrates (a-b). Consistently, the theoretical model shows that the crossover may occur at very low matrix stiffness, and thus the difference between forces generated by VIF -/- and VIF +/+ cells may become significant only at high matrix stiffness (c).

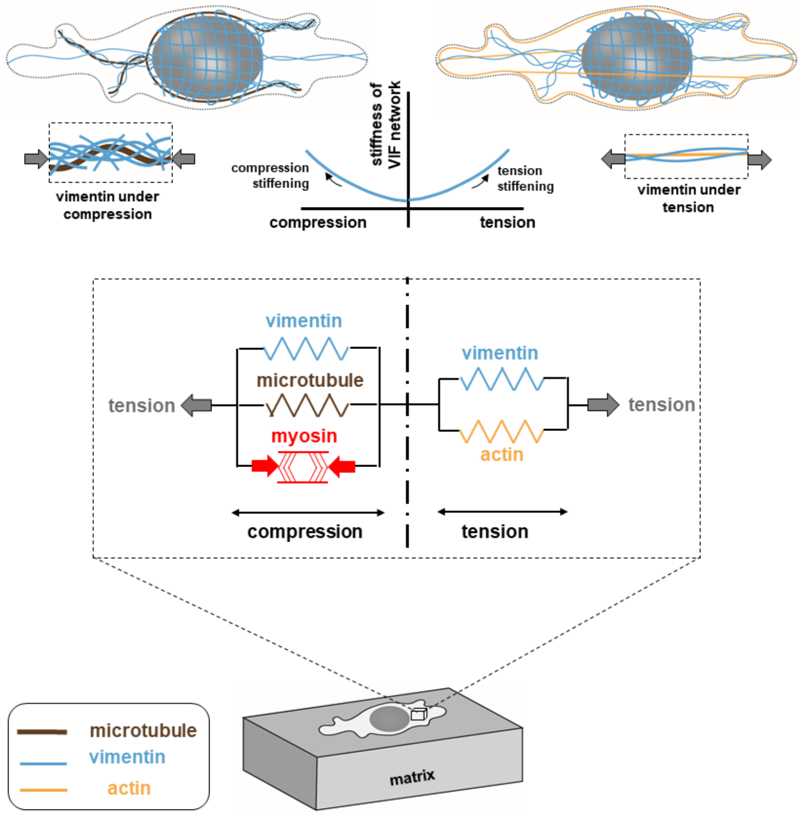


Figure S15. Vimentin intermediate filaments may experience tensile or compressive forces and they stiffen under both tension and compression. This strain stiffening can, in turn, lead to long-range force propagation in the cytoplasm as shown in Figure 4.

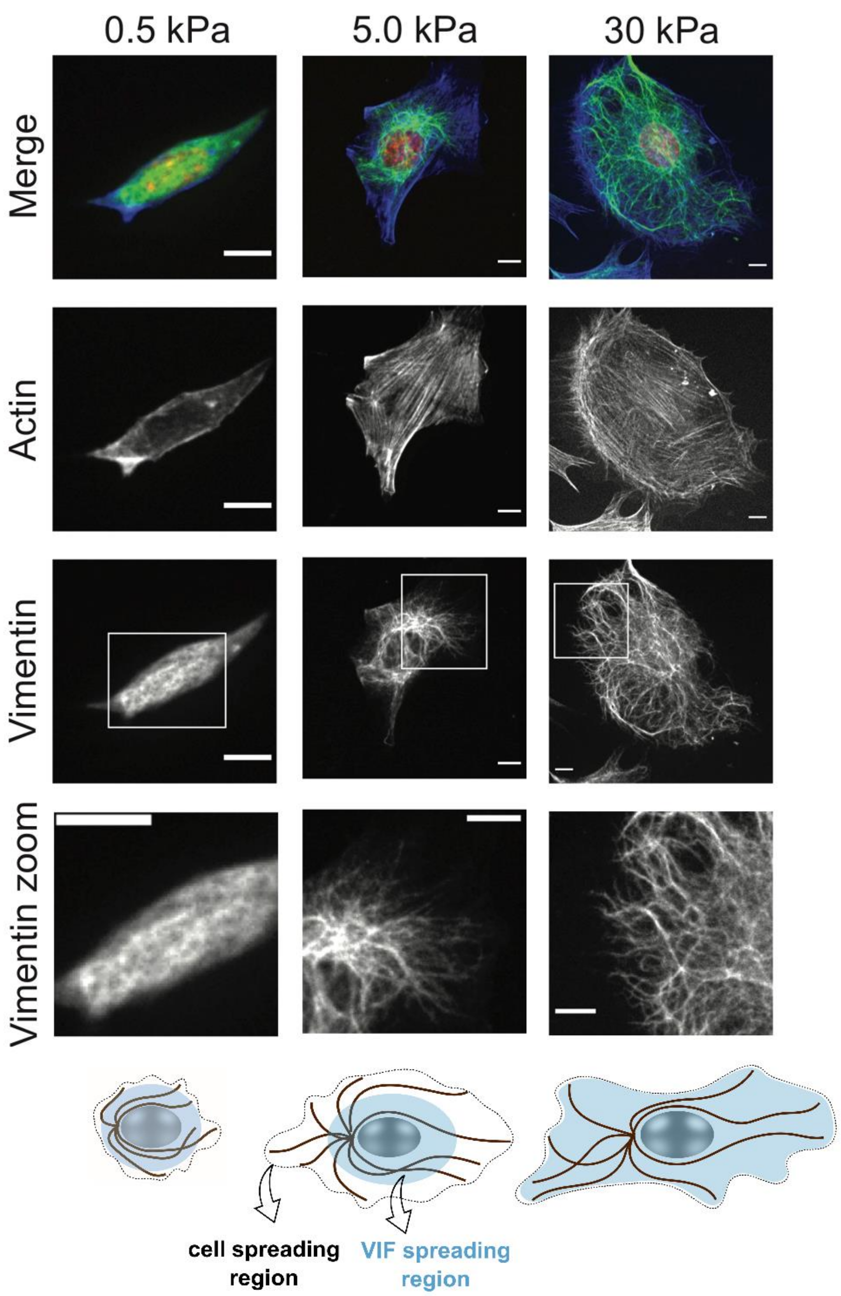


Figure S16. Fibroblasts on very soft substrates (0.5 kPa) are not able to spread. With increasing substrate stiffness, fibroblasts spread more and experience higher tension in their actin networks. Concomitant with the spreading and generation of tension, the vimentin filament network changes from a wavy mesh-like structure in the juxtanuclear region (5 kPa) to fibrous filaments which reach the cell periphery (30 kPa). Scale bars: 10 µm

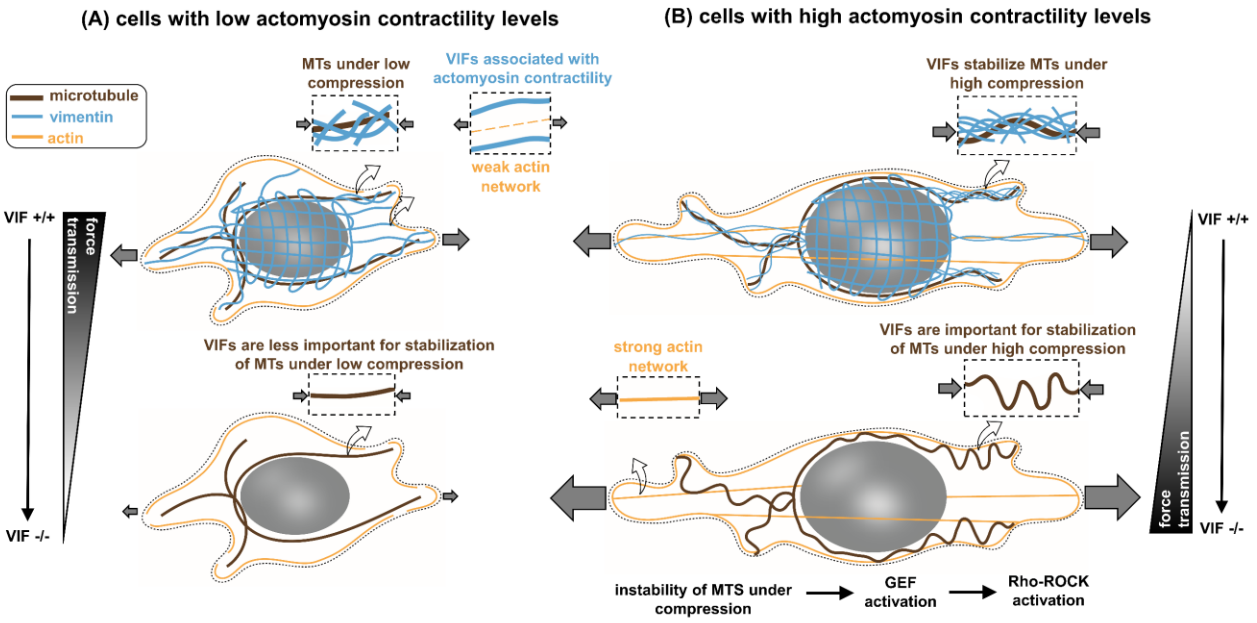


Figure S17. A model summarizing how vimentin intermediate filaments impact the transmission of mechanical forces to the extracellular matrix. Vimentin filaments undergo tensile forces and are involved in the transmission of tensile forces to the extracellular matrix (force-transmitting role). Vimentin filaments also laterally reinforce and stabilize microtubules under the contractility-based compressive forces (microtubule-reinforcing role). Therefore, disruption of vimentin can decrease (due to the force-transmitting role) or increase (due to the microtubule-reinforcing role) matrix deformation. (A) Cells with low actomyosin contractility (e.g., cells on soft matrices) experience low compression on the microtubule network, and therefore disruption of vimentin filaments does not cause significant instability in the microtubule network. Subsequently, disruption of vimentin filaments at low actomyosin regimes reduces matrix deformation as the force-transmitting role of intermediate filaments overpowers their microtubule-reinforcing role. (B) In contrast, cells with high actomyosin contractility (e.g., cells on stiff matrices) experience high compression on the microtubule network, and therefore disruption of vimentin filaments causes instability of the microtubule network which in turn can increase contractility. On the other hand, cells on stiff substrates form a strong contractile actomyosin network to transmit tensile forces to the matrix, and therefore disruption of vimentin filaments does not significantly reduce the force transmission. As a result, disruption of vimentin at high actomyosin regimes increases matrix deformation as the microtubule-reinforcing role of vimentin overpowers its force-transmitting role.

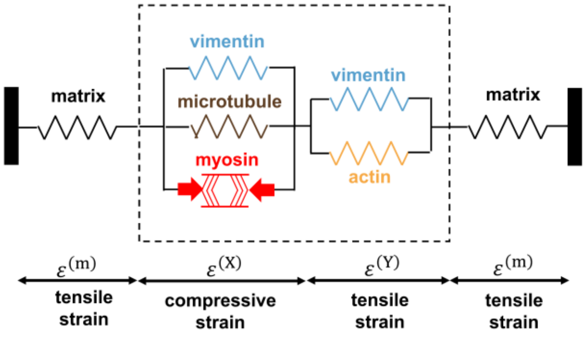


Figure S18. One-dimensional representation of the model components.

Table 1. List of parameters used in the model

| Parameter | Description | unit | value |
|---|---|---|---|
| $\alpha$ | chemo-mechanical feedback parameter | 1/(kPa) | 0.8 |
| $\beta$ | chemical stiffness parameter | 1/(kPa) | 1.2 |
| $\rho_0$ | initial contractility | kPa | 10 |
| $E^{(\mathrm{MT})}$ | elastic modulus of the microtubule network | kPa | 9 |
| $E^{(\mathrm{I})}$ | initial elastic modulus of the actin network | kPa | 0.5 |
| $m$ | actin network stiffening parameter | - | 15 |
| $\epsilon_{\mathrm{A}}$ | critical strain for the actin network stiffening | - | 0.15 |
| $E^{(\mathrm{VC})}$ | elastic modulus of the vimentin network under compression | kPa | 1.45 |

The model parameters were determined by fitting the model to Traction Force Microscopy (TFM) experiments for micropatterned fibroblasts cultured on fibronectin-coated substrates with different stiffness (2.8-30 kPa) and surface areas (700-2400 µm$^2$) (*8*). Note that $E^{(\mathrm{VC})}$ is the elastic modulus of the vimentin network under compression. The vimentin network under tesion can have the same or a different elastic modulus (we set $E^{(\mathrm{VT})}$ = 0.6 $E^{(\mathrm{VC})}$). We studied the effects of $E^{(\mathrm{VC})}$ and $E^{(\mathrm{VT})}$ on cell traction force in Figures S11 ans S12.
